# Enabling non-viral DNA delivery using lipid nanoparticles co-loaded with endogenous anti-inflammatory lipids

**DOI:** 10.1101/2024.06.11.598533

**Authors:** Manthan N. Patel, Sachchidanand Tiwari, Yufei Wang, Sarah O’Neill, Jichuan Wu, Serena Omo-Lamai, Carolann Espy, Liam S. Chase, Aparajeeta Majumdar, Evan Hoffman, Anit Shah, András Sárközy, Jeremy Katzen, Norbert Pardi, Jacob S. Brenner

**Affiliations:** Department of Systems Pharmacology and Translational Therapeutics, Perelman School of Medicine, University of Pennsylvania, Philadelphia, PA, USA; Division of Pulmonary, Allergy, and Critical Care, Department of Medicine, Perelman School of Medicine University of Pennsylvania Philadelphia, PA, USA; Department of Bioengineering, School of Engineering & Applied Science, University of Pennsylvania, Philadelphia, PA, USA; Department of Microbiology, Perelman School of Medicine, University of Pennsylvania, Philadelphia, PA, USA

## Abstract

Lipid nanoparticles (LNPs) have transformed genetic medicine, recently shown by their use in COVID-19 mRNA vaccines. While loading LNPs with mRNA has many uses, loading DNA would provide additional advantages such as long-term expression and availability of promoter sequences. However, here we show that plasmid DNA (pDNA) delivery via LNPs (pDNA-LNPs) induces acute inflammation in naïve mice which we find is primarily driven by the cGAS-STING pathway. Inspired by DNA viruses that inhibit this pathway for replication, we co-loaded endogenous lipids that inhibit STING into pDNA-LNPs. Specifically, loading nitro-oleic acid (NOA) into pDNA-LNPs (NOA-pDNA-LNPs) ameliorates serious inflammatory responses *in vivo* enabling prolonged transgene expression (at least 1 month). Additionally, we demonstrate the ability to iteratively optimize NOA-pDNA-LNPs’ expression by performing a small LNP formulation screen, driving up expression 50-fold *in vitro*. Thus, NOA-pDNA-LNPs, and pDNA-LNPs co-loaded with other bioactive molecules, will provide a major new tool in the genetic medicine toolbox, leveraging the power of DNA’s long-term and promoter-controlled expression.

## Introduction

The success of the COVID-19 vaccines showed the unprecedented power of lipid nanoparticles (LNPs) to deliver nucleic acids to target cells, driving levels of expression of encoded proteins higher than prior non-viral technologies^1^. This triumph spurred the biopharma industry to invest billions of dollars in LNP-based therapeutics, where LNPs’ ability to be targeted to specific organs and cell types has enabled applications in many types of disease^2^. In the COVID-19 vaccines and many other applications of LNPs, the nucleic acid cargo has been mRNA, which can transiently express encoded proteins. However, LNPs’ sole focus on mRNA delivery may prevent LNPs from reaching their full therapeutic potential, as mRNA has a relatively short half-life (∼hours), lacks a promoter region to achieve cell-type-specific and temporal control, and is not stable for long at room temperature or 4°C^3^.

DNA could overcome many of the challenges associated with the use of mRNA, and thereby open up new applications for LNPs. DNA can express transgene proteins for several months^4^, has a promoter that can be made cell-type-specific and/or turned on/off with small molecule drugs (such as a doxycycline-sensitive promoter)^5^, and can be stored for months at 4°C. Such advantages of DNA could open up LNPs’ applications to include long-term expression, including monoclonal antibodies, secreted proteins, or even intracellular proteins, with the minimal concerns about the half-life of the engineered proteins because of constant protein production. Additionally, DNA can be used to express short-hairpin RNA (shRNA) to knockdown proteins long-term, gene editing proteins and guide RNAs.

For each of these genetic cargo, DNA-loaded LNPs (DNA-LNPs) would offer the advantages of long-term expression (and thus infrequent dosing) and low cost-of-goods-sold (COGS), along with the advantages LNPs already provide, including high levels of expression, low immunogenicity (compared to viral vectors)^6^, and fewer limitations on cargo size^7^. These collective benefits of DNA-LNPs would enable treatment of diseases that are less accessible to mRNA-LNPs, such as diseases of chronic autoimmunity, degeneration, pain, and more.

While DNA seems like a natural fit for LNPs, DNA-LNPs have witnessed very few publications in the last 15-20 years since LNPs were developed. Here, we begin by showing that this lack of study is likely because plasmid DNA delivered via LNPs (pDNA-LNPs) is highly inflammatory and induces mortality at commonly used therapeutic doses in naïve mice. Through *in vitro* immunostaining and *in vivo* knockout mouse model, we found that this inflammation is largely driven by activation of the cGAS-STING signaling pathway, comprised of cyclic GMP-AMP synthase (cGAS) and stimulator of interferon genes (STING). cGAS is a cytosolic DNA sensor that recognizes DNA via electrostatic interactions^8^. Upon DNA binding, cGAS activates downstream STING signaling which leads to massive inflammation characterized by upregulation of type 1 interferons (IFNs), such as IFN-β, and pro-inflammatory cytokines, such as IL-6^9^. As cGAS can detect DNA in a sequence-independent manner, we pursued an alternative strategy to mitigate pDNA-induced inflammation: co-loading naturally occurring STING inhibitors into pDNA-LNPs to enable safe and effective delivery.

We co-loaded nitro-oleic acid (NOA), an endogenous anti-inflammatory lipid that inhibits STING^10^, into pDNA-LNPs (NOA-pDNA-LNPs). This led to undetectable levels of pDNA-induced STING activation *in vitro*. While 1 mg/kg (∼25 μg) of standard pDNA-LNPs injected intravenously (IV) into naïve C57BL/6 mice induced 100% mortality within 2 days, NOA-pDNA-LNPs induced 0% mortality. We then showed that addition of NOA in pDNA-LNPs does not hinder transgene expression and enables prolonged protein expression (at least 1 month). Additionally, we showed the potential to further optimize the nascent technology of pDNA-LNPs, as a small LNP formulation screen improved the transgene expression capacity of NOA-pDNA-LNPs 50-fold *in vitro*, enabling NOA-pDNA-LNPs to achieve similar transfection efficiencies to one of the gold-standard *in-vitro* transfection reagents, Lipofectamine, in a cell type that is widely considered difficult-to-transfect, human induced-pluripotent-stem-cell (iPSC)-derived type II alveolar epithelial cells (iAT2s).

Overall, these results show that NOA-pDNA-LNPs, and more generally pDNA-LNPs co-loaded with bioactive molecules, have the potential to provide safe, long-term expression of therapeutic cargo.

## Results

### Unlike nucleoside-modified mRNA-LNPs, pDNA-LNPs induce serious inflammatory responses in wild type mice

To systematically probe inflammation, we used BioNTech/Pfizer’s FDA-approved COVID-19 mRNA vaccine LNP formulation, containing the ionizable lipid ALC-0315, formulated with either 5moU nucleoside-modified mRNA or plasmid DNA (pDNA). To minimize carrier side effects as a confounding variable, both mRNA- and pDNA-LNPs were formulated using 40-to-1 total lipid-to-nucleic acid ratio (w/w).

After IV injecting a commonly used therapeutic dose^11^ of 1 mg/kg (∼25 μg pDNA per mouse) into naïve C57BL/6 mice, we observed 100% mortality within 2 days for mice that received pDNA-LNPs, compared to 0% for ones that received mRNA-LNPs (**Fig. 1a**). Furthermore, we noticed extreme lethargy and lack of movement of mice 4-hours after pDNA-LNP administration, using an artificial intelligence (AI) motion-tracking device that we previously validated for detecting infusion reactions^12^ (**Fig. 1b**). Thus, LNPs delivering pDNA at a 1 mg/kg dose, but not nucleoside-modified mRNA at the same dose, induce an acute inflammatory reaction that leads to death within 2 days in naïve mice.

**Figure 1:**
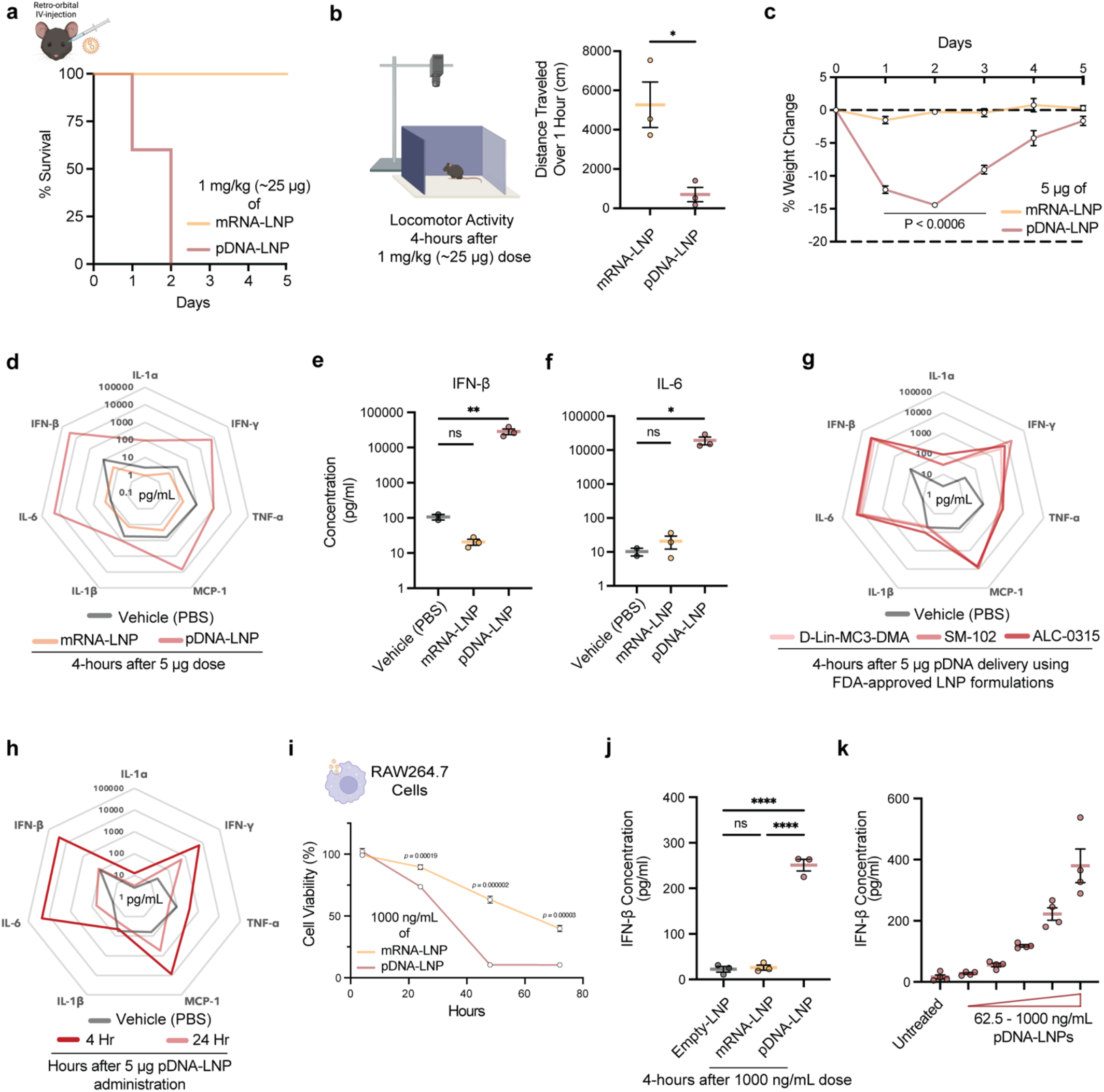
Unlike nucleoside-modified mRNA-LNP, pDNA-LNP delivery causes acute inflammatory responses. **a**, Survival curve graph of naïve C57BL/6 (“Black-6”) mice IV injected with 1 mg/kg of either mRNA- or pDNA-LNP shows 100% mortality in mice treated with pDNA-LNPs compared to 0% in mice treated with mRNA-LNPs. **b**, 4-hours after 1 mg/kg IV injection of LNPs, movement of mice was tracked for 1 hour and total distance walked was assessed by AI software (DeepLabCuts), a validated metric of an infusion reaction. Mice treated with pDNA-LNPs have significantly lower total distance walked compared to mRNA-LNP control, indicating severe lethargy. **c**, Weight change over time in mice given a much lower IV dose (5 μg) of mRNA- or pDNA-LNPs. **d, e, f**, Multiplex analysis of pro-inflammatory plasma cytokines 4-hours post 5 μg IV dose of pDNA-LNP indicates acute systemic inflammation compared to PBS and mRNA-LNP controls. Specifically, IFN-β (**e**) and IL-6 (**f**) levels are ∼1400-fold and ∼1000-fold higher, respectively, for mice injected with pDNA-LNP compared to mRNA-LNP. **g**, 5 μg of pDNA formulated with three FDA-approved LNP formulations (D-Lin-MC3-DMA, SM-102, and ALC-0315 ionizable lipids) were IV injected and plasma was collected 4-hours later for cytokine quantification, which indicated this inflammatory response occurs across various LNP formulations. **h**, Cytokine levels in mouse plasma collected 4- or 24-hours after IV injection of 5 μg of pDNA-LNPs, highlighting acute-but-transient inflammation, with the majority of pro-inflammatory cytokine levels back to baseline at the 24-hour time point. **i – k**, *In vitro* studies in a macrophage-derived cell line, RAW264.7. **i**, Effect of 1000 ng/mL mRNA- or pDNA-LNP on cell viability over time. **j**, IFN-β levels in cell supernatant 4-hours after exposure to 1000 ng/mL of empty-, mRNA-, and pDNA-LNP show DNA-cargo-specific inflammatory cytokine production. **k**, IFN-β levels in cell supernatant increase exponentially as a function of pDNA-LNP dose. ***Statistics***: n=5/group for **a** and **c** (biological replicates), n=4/group for **k** (biological replicates), n=3/group for rest (biological replicates). Data shown represents mean ± SEM. **b, c, i**, Unpaired t-tests were performed. For all other graphs, comparisons were made using one-way ANOVA with Tukey’s post-hoc test.

To study the mechanisms underlying pDNA-LNP-induced inflammation, we reduced the pDNA-LNP dose 5-fold to 5 μg pDNA per mouse, ensuring survival and enabling assessment of signaling pathways. Just one day after the administration of 5 μg pDNA-LNP, the mice lost >10% of their body weight, taking ∼5 days to return to baseline levels (**Fig. 1c**). Furthermore, to assess systemic inflammation, we collected plasma 4-hours after the pDNA-LNP IV injection and observed a significant increase in the levels of various pro-inflammatory cytokines (**Fig. 1d**). Specifically, IFN-β and IL-6 plasma concentrations increased 1400- and 1000-fold, respectively, compared to the mRNA-LNP-treated group (**Fig. 1e, 1f**). Thus, LNPs loaded with pDNA, but not with nucleoside-modified mRNA, induce a massive cytokine response. Notably, IFN-β is a type 1 IFN that is traditionally known as an antiviral cytokine^13^, hinting that the immune system likely identifies pDNA-LNP administration as a viral infection.

To ensure pDNA-LNP toxicity was independent of LNP formulation, we synthesized pDNA-LNPs using 2 other FDA-approved LNP formulations (D-Lin-MC3-DMA and SM-102 ionizable lipids)^14,15^. Regardless of LNP formulation, we observed acute inflammation *in vivo* (**Fig. 1g**). Moreover, we observed the same inflammatory response in LNPs loaded with various plasmids (**Supplementary Fig. 1**).

We next investigated the kinetics of the inflammatory response, as the mice returned to a normal visual and behavioral phenotype 24-hours after pDNA-LNP administration. In agreement with our visual assessment, we observed a decrease in most pro-inflammatory cytokines in the plasma 24-hours after the 5 μg pDNA-LNP dose, with the majority of cytokines – specifically IFN-β and IL-6 – back to baseline levels (**Fig. 1h**). Furthermore, we ensured inflammation does not reappear at later time points by measuring plasma cytokines 5-days after pDNA-LNP administration (**Supplementary Fig. 2**). Since this response was acute, we investigated if pDNA-LNPs are inflammatory in innate immune cells such as macrophages that we previously showed are key in mediating acute LNP carrier-related toxicities and is known to sensitively detect DNA-viruses^16,17^.

**Figure 2:**
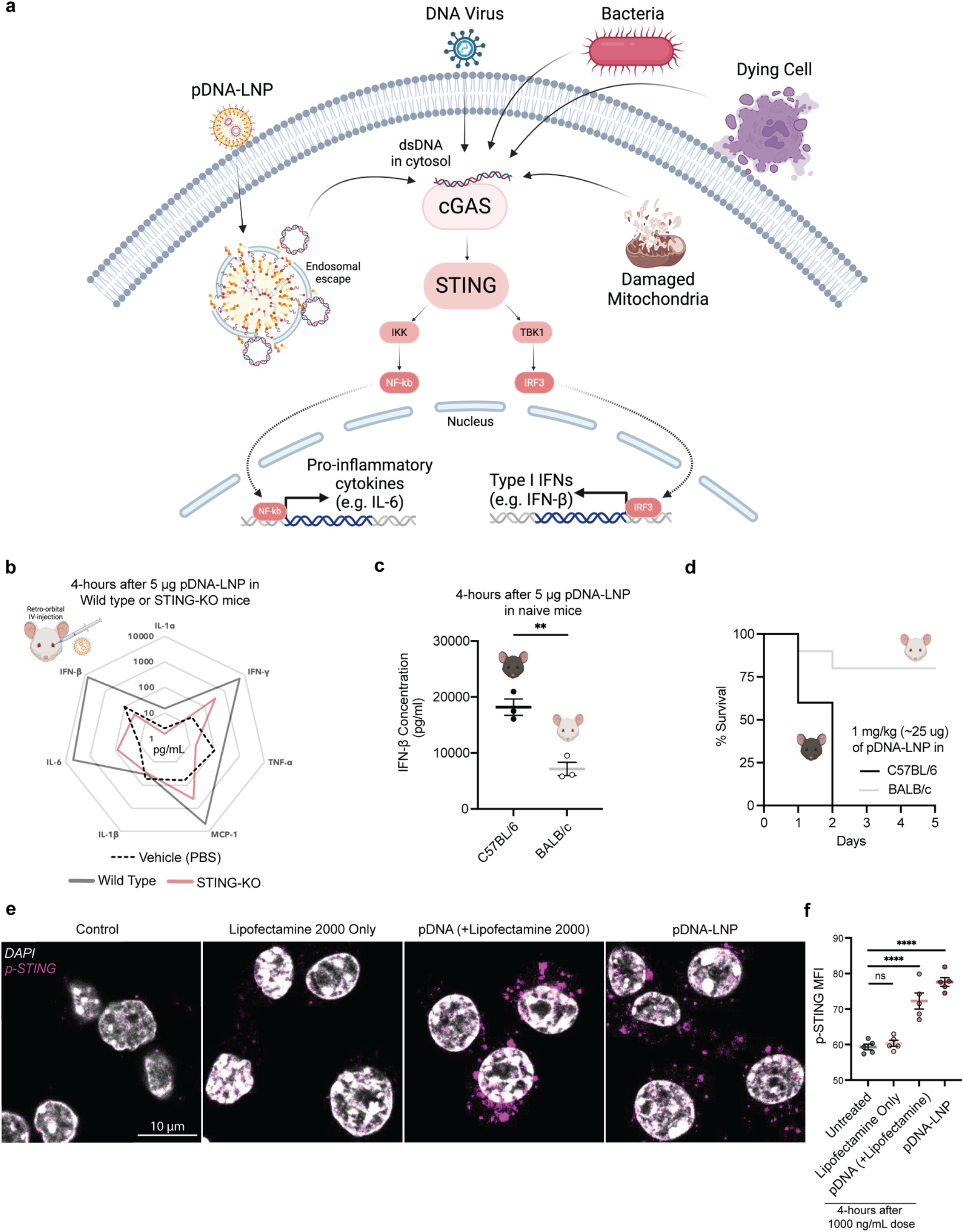
STING activation drives pDNA-LNP-induced inflammation. **a**, The proposed mechanism that drives pDNA-LNP inflammation. Any cytosolic DNA (endogenous or exogenous) is detected - independent of DNA sequence - by cGAS, leading to downstream activation of STING which induces an acute inflammatory response (modified from). **b**, STING knockout (STING-KO) mice IV injected with 5 μg pDNA-LNPs have reduced levels of pro-inflammatory cytokines in plasma 4-hours after dose compared to wild type BALB/c. **c, d**, pDNA-LNPs injected in C57bl/6 (“Black-6”) and BALB/c have varying levels of inflammation with ∼2-fold lower IFN-β levels in BALB/c mice leading to improved survival rates at 1 mg/kg dose (**d**). **e, f**, Representative images of phosphorylated STING (p-STING) 4-hours post treatment indicates STING activation for pDNA group (1000 ng/mL dose) in RAW264.7 cells (**e**) Quantification of p-STING mean fluorescence intensity (MFI) from original images (**f**). ***Statistics***: n=3/group for **b** and **c** (biological replicates), n=5/group for **d** (biological replicates), n=5/group for **f** (technical replicates, similar findings with biological replicates). Data shown represents mean ± SEM. **c**, Unpaired t-test was performed. **f**, Comparisons were made using one-way ANOVA with Tukey’s post-hoc test.

We treated the macrophage-derived cell line, RAW264.7, with various concentrations of pDNA-LNPs, for various treatment times, and measured cell viability (**Supplementary Fig. 3**). At a concentration of 1000 ng/mL for both mRNA- and pDNA-LNPs, we found a significant decrease in cell viability over time, with ∼90% cell death 48-hours after pDNA-LNP compared to <50% after mRNA-LNP treatment (**Fig. 1i**). Moreover, similarly to the *in vivo* cytokine results, we observed a significant increase (∼10-fold) in IFN-β levels in the cell supernatant for pDNA-LNPs compared to empty- and mRNA-LNP controls 4-hours after 1000 ng/mL LNP treatment (**Fig. 1j**). Lastly, we performed a dose response study of pDNA-LNPs which showed exponential increase in IFN-β levels (**Fig. 1k**).

**Figure 3:**
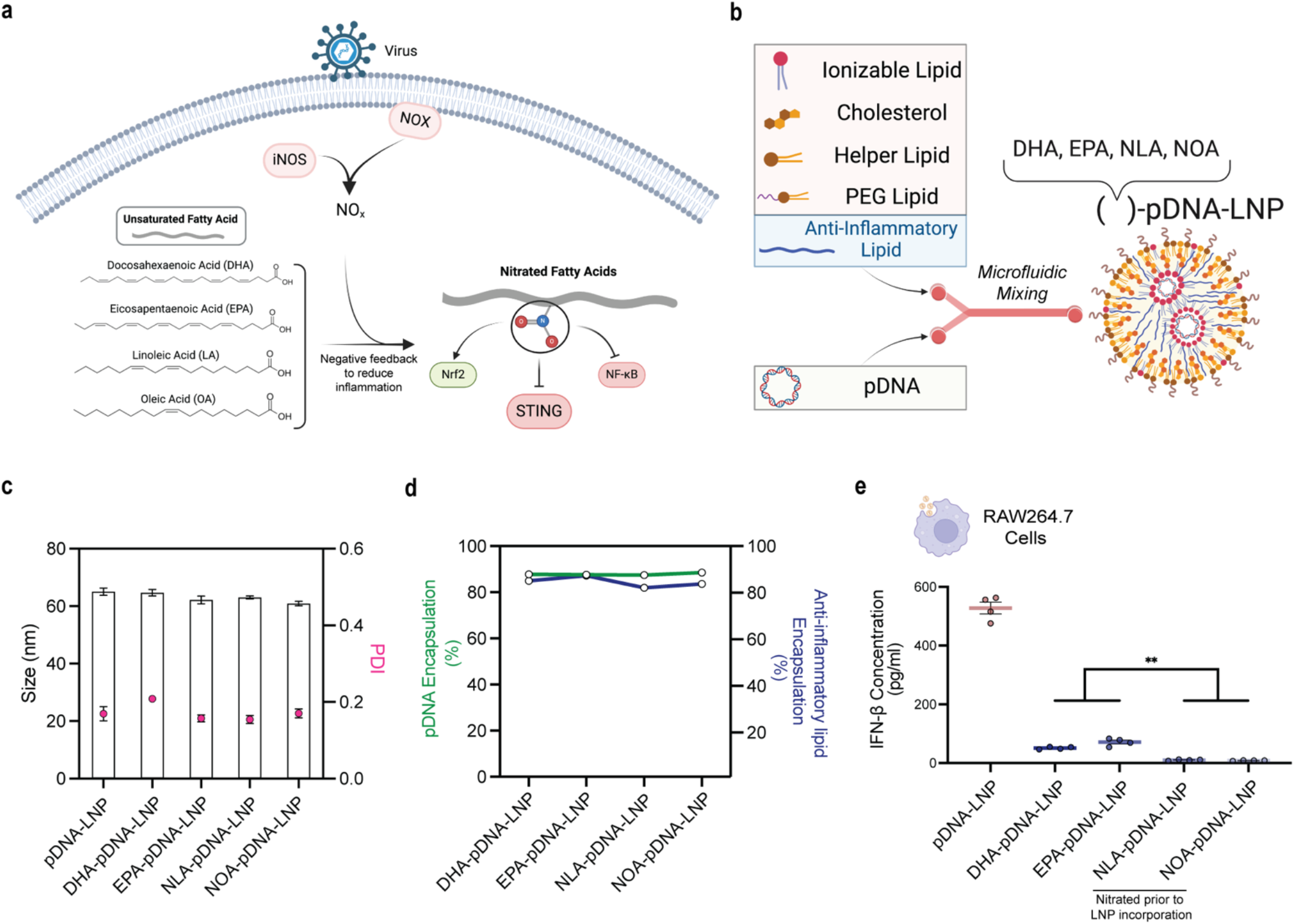
Co-loading endogenous anti-inflammatory lipids with STING inhibitory activity into standard pDNA-LNP formulations. **a**, Schematic showing how cell stress caused by virus infections leads to nitration of endogenous unsaturated fatty acids to form nitrated fatty acids that have anti-inflammatory properties, including potent inhibition of STING (modified from Fig. 1 of the published paper by Hansen et al., PNAS 2018). **b**, Development of platform technology by adding anti-inflammatory lipids (AILs) as the 5^th^ component in the lipid mixture prior to LNP formation using microfluidics. **c**, Size distribution of pDNA-LNPs loaded with various AILs determined by dynamic light scattering (DLS) shows no differences in LNP size or polydispersity index (PDI). **d**, All tested AILs load >80% (AIL-to-lipid ratio of 0.2, mole-to-mole) and have no impact on pDNA encapsulation. **e**, pDNA-LNPs loaded with various AILs reduce IFN-β levels in cell supernatant 4-hours after 1000 ng/mL dose. AILs that are nitrated prior to LNP formation (NLA and NOA) are more effective than lipids that are not nitrated during LNP formation (DHA and EPA). ***Statistics*:** n=4/group for **e** (biological replicates), n=3/group for **c** (technical replicates, similar findings with biological replicates). Data shown represents mean ± SEM and comparisons were made using one-way ANOVA with Tukey’s post-hoc test.

Prior to investigating the mechanisms that mediated pDNA-LNP-induced toxicities, we inquired if the route of administration led to varying levels of inflammation. We administered mRNA- and pDNA-LNPs intratracheally (to the lungs) at a dose of 5 μg. We evaluated inflammation specific to the lungs by examining protein and leukocyte levels in the bronchoalveolar lavage (BAL) fluid, which indicates capillary leakage and leukocyte penetration into the alveoli (air sacs). We observed higher levels of protein, total cells, and pro-inflammatory cytokines in the BAL from mice that received pDNA-LNPs compared to mRNA-LNPs, highlighting pDNA’s cargo-induced inflammatory responses (**Supplementary Fig. 4**).

**Figure 4:**
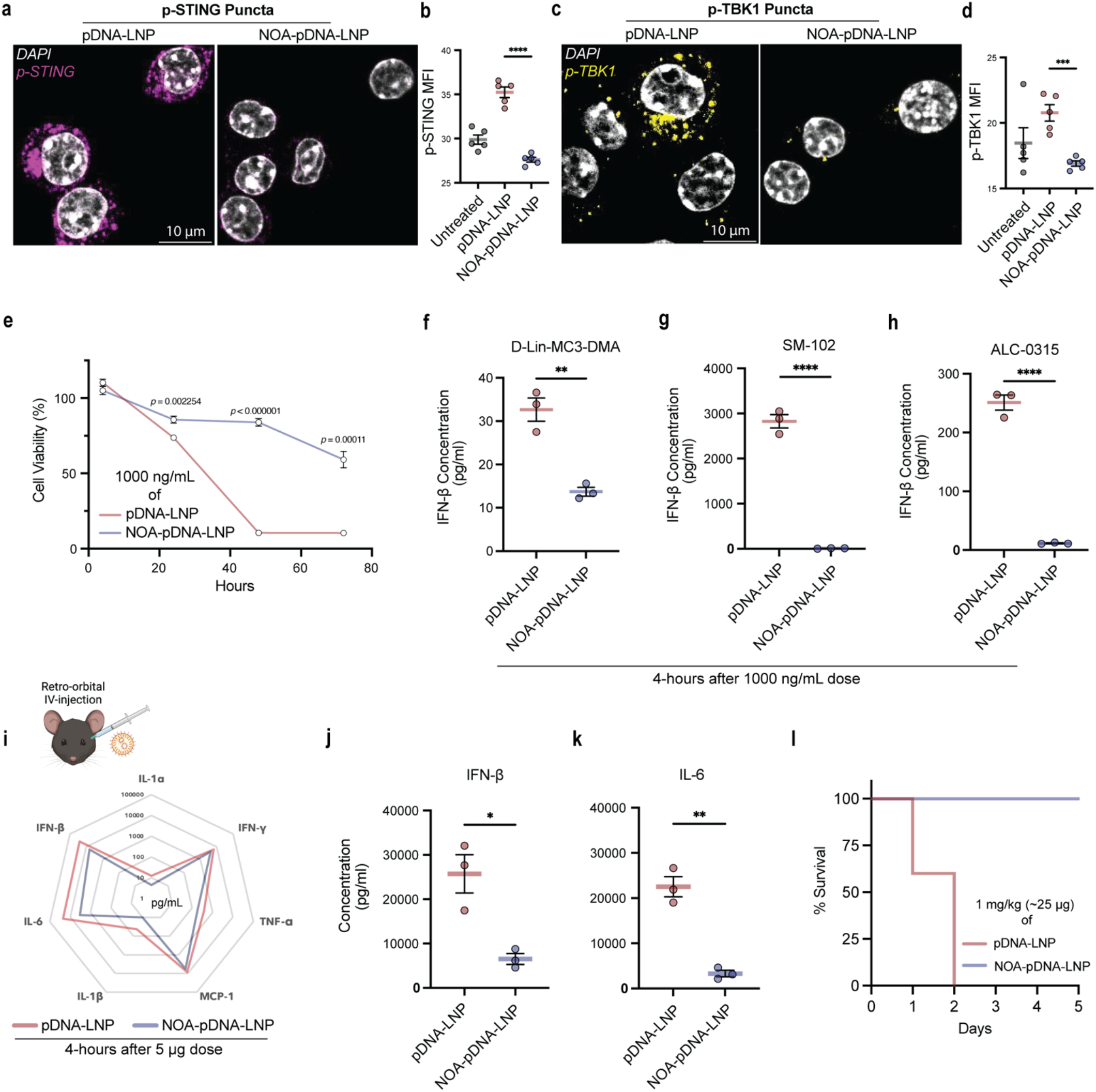
Nitro-oleic acid (NOA) loaded pDNA-LNPs (NOA-pDNA-LNPs) show superior safety profiles *in vitro* and *in vivo*. **a, b**, Representative confocal images of phosphorylated STING (p-STING) show NOA-pDNA-LNPs do not activate STING compared to standard pDNA-LNPs in RAW264.7 cells. Quantification of p-STING mean fluorescence intensity (MFI) shows significant decrease for NOA-pDNA-LNP group relative to standard pDNA-LNP (**b**). **c, d**, Similar to (**a**) and (**b**), confocal imaging of downstream marker of STING activation, phosphorylated TBK1 (p-TBK1), is also not activated for cells treated with NOA-pDNA-LNPs compared to standard pDNA-LNPs. **e**, RAW264.7 cell viability measured over time indicates better tolerability of NOA-pDNA-LNPs compared to standard pDNA-LNPs. **f, g, h**, IFN-β levels in cell supernatant are significantly lower for NOA-pDNA-LNPs compared to standard pDNA-LNP 4-hours post 1000 ng/mL dose, irrespective of LNP formulation [(**f**) D-Lin-MC3-DMA, (**g**) SM-102, and (**h**) ALC-0315: all FDA-approved LNP formulations]. **i, j, k**, Quantification of pro-inflammatory plasma cytokines 4-hours post 5 μg dose of pDNA- or NOA-pDNA-LNPs. Specifically, IFN-β (**j**) and IL-6 (**k**) levels are ∼4-fold and ∼8-fold lower, respectively, for mice injected IV with NOA-pDNA-LNP compared to standard pDNA-LNP control. **l**, Survival curve in C57BL/6 (“Black-6”) mice, comparing IV dose of 1 mg/kg of pDNA-LNP and NOA-pDNA-LNPs shows addition of NOA in pDNA-LNPs completely prevents mortality. ***Statistics*:** n=5/group for **b** and **d** (technical replicates, similar findings with biological replicates), n=3/group for **e-k** (biological replicates), n=5/group for **l** (biological replicates). Data shown represents mean ± SEM. **e-h, j, k**, Unpaired t-tests were performed. **b, d**, Comparisons were made using one-way ANOVA with Tukey’s post-hoc test.

In summary, we show that pDNA-LNPs induce acute inflammatory responses in naïve C57BL/6 mice that results in mortality when IV injected at a commonly used therapeutic dose of 1 mg/kg.

### STING activation drives pDNA-LNP-induced inflammation

Amongst the well-characterized DNA-sensing pathways, cGAS-STING was frequently reported to induce acute inflammatory responses and significantly upregulate type 1 IFNs in response to viral infections^8,9,13^. Interestingly, cGAS activation can occur not only from viral DNA, but any cytosolic DNA^18^. Thus, it serves as a versatile sensor that can also be activated by bacterial DNA, DNA from dying cells, and even self-mitochondrial DNA that is released into the cytosol during cell stress leading to downstream activation of STING (**Fig. 2a**). As such, we investigated if pDNA-LNPs activate this pathway.

To assess the role of the cGAS-STING pathway in pDNA-LNP-induced inflammation, we IV injected pDNA-LNPs in wild type and STING knock out (STING-KO) mice and collected plasma 4-hours later. Note that we used BALB/c mice instead of C57BL/6 mice as the only STING-KO mice that were commercially available were of the BALB/c strain. After IV injection of 5 μg of pDNA-LNPs, we observed significantly lower levels of various pro-inflammatory cytokines in STING-KO mice compared to wild type mice, indicating STING activation as a primary driver of pDNA-LNP inflammation (**Fig. 2b**). Importantly, IFN-β and IL-6 were down to baseline levels in the STING-KO mice injected with pDNA-LNPs.

Furthermore, we observed the overall level of inflammation was lower in the male BALB/c strain compared to male C57BL/6 strain (∼2-fold lower IFN-β levels), though both are much above the baseline and the mRNA-LNP control (**Fig. 2c**). Additionally, we found that BALB/c mice better tolerated pDNA-LNPs as the mortality rate was 20% compared to 100% in C57BL/6 mice at the 1 mg/kg (∼25 μg) dose (**Fig. 2d**). Notably, BALB/c mice that survived still experienced significant weight loss (**Supplementary Fig. 5**).

**Figure 5:**
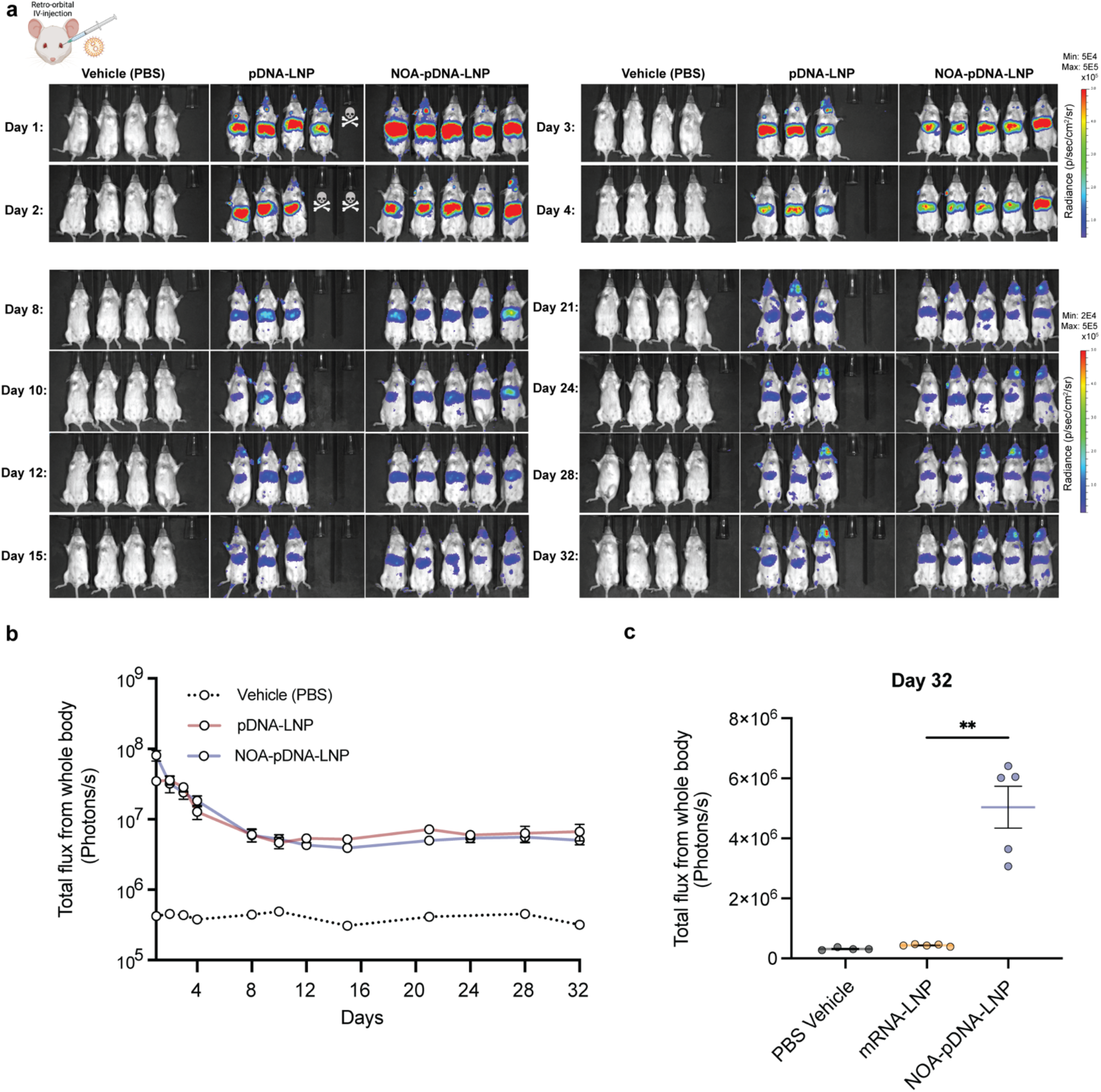
NOA-pDNA-LNPs show prolonged transgene expression *in vivo*. **a**, IVIS images of BALB/c mice that were IV injected (retro-orbitally) with 25 μg of either pDNA-LNP or NOA-pDNA-LNPs (encoding luciferase). **b**, Quantified total flux (photons/s) from IVIS images show addition of NOA does not hinder pDNA transgene expression capacity and shows prolonged expression (at least 1 month). **c**, At day 32, total transgene expression levels are significantly higher in mice treated with 25 μg NOA-pDNA-LNPs compared to PBS and mRNA-LNP (5 μg) controls. ***Statistics***: n=4/group for PBS control group and n=5/group for pDNA-LNP and NOA-pDNA-LNP groups for **a-c** (biological replicates). Note that two mice that received 25 μg of pDNA-LNPs died within 2-days. Data shown represents mean ± SEM. **c**, Comparisons were made using one-way ANOVA with Tukey’s post-hoc test.

In addition, using confocal microscopy, we confirmed STING activation in macrophages treated with pDNA - using lipofectamine or LNP - by staining for activated STING (phosphorylated-STING or p-STING) (**Fig. 2e**). Note that we do not see STING activation in the lipofectamine-only control, indicating it is a pDNA-specific activation rather than carrier-based activation (**Fig. 2f**).

### Developing a platform technology: co-loading of endogenous anti-inflammatory lipids with STING inhibitory activity into standard pDNA-LNP formulations

We next looked into inhibitors of STING that we can co-load into standard pDNA-LNP to ameliorate its acute adverse events. We first investigated anti-inflammatory lipids (AILs) as they are highly lipophilic and have better chance of loading into standard pDNA-LNPs compared to small molecule drugs. We also prioritized fast-acting drugs as opposed to siRNAs, since pDNA-LNP inflammation is acute-but-transient, as shown previously in **Fig. 1h**.

Interestingly, many unsaturated fatty acids - such as Docosahexaenoic acid (DHA), Eicosapentaenoic acid (EPA), Linoleic acid (LA), and Oleic acid (OA) - have been found to be nitrated after a virus infection as a negative feedback loop to dampen excessive inflammation^10,19^ (**Fig. 3a**). These nitrated fatty acids (NFAs) act as electrophiles with the ability to modify various proteins on specific exposed cysteines, leading to inhibition of nuclear factor kappa B (NF-κB), which controls an array of prototypical pro-inflammatory signaling genes, and activation of nuclear factor-erythroid 2 related factor 2 (Nrf2), which controls an array of antioxidant response element–dependent genes^20^. Most importantly, NFAs are potent inhibitors of STING^10^.

We loaded DHA, EPA, and already nitrated versions of LA and OA (NLA and NOA, respectively) by adding these AILs as a fifth component into standard LNP formulation at an AIL-to-total lipids ratio of 0.2 (mole-to-mole) (**Fig. 3b**). Using dynamic light scattering, we confirmed LNP size and polydispersity were unaffected by the addition of this fifth component (**Fig. 3c**). Furthermore, all AILs showed encapsulation efficiencies of >80% without negatively affecting pDNA loading (**Fig. 3d**).

To initially assess the safety profile of AIL-loaded pDNA-LNPs, we treated RAW264.7 cells with a 1000 ng/mL dose and quantified IFN-β levels in the cell supernatant 4-hours after treatment. All AIL-loaded pDNA-LNPs had significantly reduced IFN-β levels compared to standard pDNA-LNPs (**Fig. 3e**). Furthermore, pDNA-LNPs loaded with NLA or NOA (lipids nitrated prior to LNP incorporation) performed better compared to pDNA-LNPs loaded with DHA or EPA. Note that we did not observe a decrease in cell viability of NOA-pDNA-LNPs 4-hours after administrating at 1000 ng/mL dose (**Supplementary Fig. 6**).

**Figure 6:**
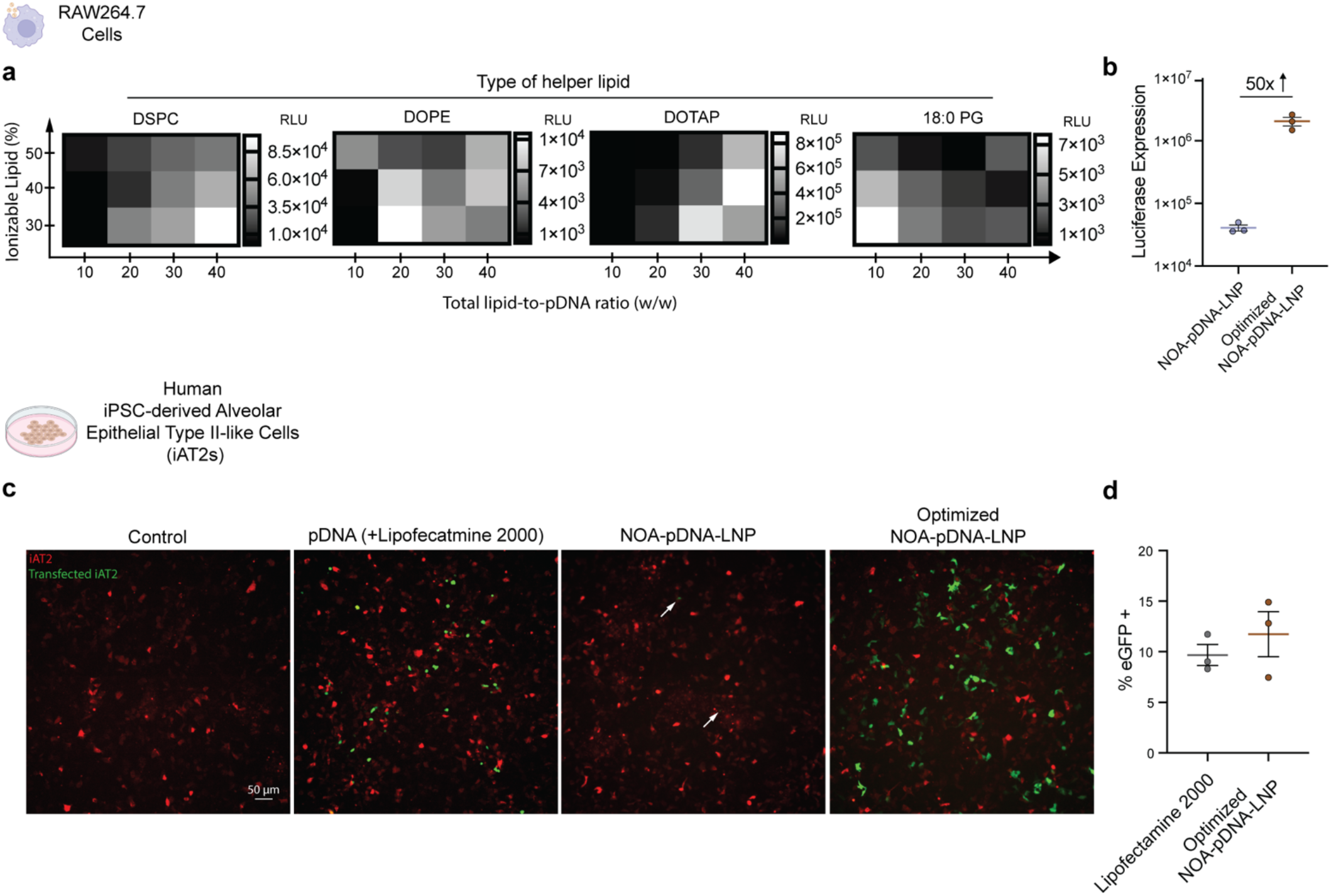
A small LNP formulation screen significantly boosts transgene expression of NOA-pDNA-LNPs *in vitro*, enabling efficient transfection in difficult-to-transfect cells. **a, b**, Design of experiment (DoE) screening to optimize NOA-pDNA-LNP formulation in RAW264.7 cells. Using JMP software, a full factorial screen was designed by varying ionizable lipid mol% (30 to 50), total lipid to pDNA w/w ratio (10:1 to 40:1), and the type of helper lipid used (DSPC, DOPE, DOTAP, and 18:0 PG). All LNPs contained 0.2 D/L of NOA. 24-hour after 500 ng/mL treatment in RAW264.7 cells, luciferase protein expression was measured highlighting the influence of LNP formation parameters for transgene expression (**a**). NOA-pDNA-LNP optimized with DOTAP as the helper lipid led to 50-fold increase in transgene expression, when compared to standard NOA-pDNA-LNP (**b**). **c, d**. 2D monoculture of difficult-to-transfect cell line, human induced-pluripotent-stem-cell-derived type II alveolar epithelial cells (iAT2s), were treated with 1000 ng of eGFP encoding pDNA - using lipofectamine, standard NOA-pDNA-LNPs, or optimized NOA-pDNA-LNPs - and imaged after 48-hours. tdTomato (red) signal indicates iAT2 positively, while eGFP (green) signal indicates successfully transfected cells (**c**). 120-hours post transfection, tdTomato + cells that were also eGFP + were quantified using flow cytometry indicating similar transfection levels (trending higher) for optimized NOA-pDNA-LNPs to the gold standard, lipofectamine 2000 (**d**). ***Statistics*:** n=2 per LNP formulation for **a** (biological replicates), n=3/group for **b** and **d** (biological replicates). Data shown represents mean ± SEM.

We proceeded to use NOA for rest of the studies as it was more anti-inflammatory than DHA and EPA and more cost-effective than NLA.

### NOA-loaded pDNA-LNPs (NOA-pDNA-LNPs) show superior safety profiles *in vitro* and *in vivo*

To confirm the reduction in IFN-β production of macrophages treated with NOA-pDNA-LNPs is due to STING inhibition, we performed confocal imaging, staining for activated STING (phosphorylated-STING or p-STING). 4-hours after 1000 ng/mL dose, there was no measurable STING activation in cells treated with NOA-pDNA-LNPs (**Fig. 4a, 4b**). We also did not detect activation of TANK-binding Kinase 1 (TBK1), a downstream marker of STING activation, in cells that received NOA-pDNA-LNPs (**Fig. 4c, 4d**).

Furthermore, likely due to the reduced levels of pro-inflammatory cytokines, NOA-pDNA-LNPs maintained higher cell viability over time compared to standard pDNA-LNPs highlighting the protective effects of NOA *in vitro* (**Fig. 4e**). To generalize NOA’s anti-inflammatory effects, we loaded NOA into three different LNPs formulated with the same components found in the FDA-approved LNPs (D-Lin-MC3-DMA, SM-102, and ALC-0315). Regardless of the LNP composition, addition of NOA leads to a significant reduction in IFN-β secretion in cell supernatant 4-hours after 1000 ng/mL dose (**Fig. 4f, 4g, 4h**). Notably, the total levels of IFN-β vary with each ionizable lipid used (SM-102 > ALC-0315 > D-Lin-MC3-DMA) which we found also correlated with the transgene expression (**Supplementary Fig. 7**).

Next, we investigated the safety profile of NOA-pDNA-LNPs in naïve C57BL/6 mice. To assess the effect on acute response to pDNA, we IV injected 5 μg of standard pDNA-LNP or NOA-pDNA-LNP in mice and collected plasma 4-hours later for cytokine analysis. We observed significantly lower levels of various pro-inflammatory cytokines (IFN-β, IL-1α, IFN-γ, TNF-α, MCP-1, IL-1β, and IL-6) in mice treated with NOA-pDNA-LNPs compared to pDNA-LNPs (**Fig. 4i**). Specifically, IFN-β and IL-6 were reduced ∼4-fold and ∼8-fold, respectively (**Fig. 4j, 4k**). By performing a dose-response study, we identified NOA-to-total lipids ratio (mole-to-mole) of 0.2-0.8 as the optimal formulation parameter in ameliorating pDNA-induced inflammation (**Supplementary Fig. 8**).

We attempted to further reduce the pDNA-induced inflammatory response by loading A151, an oligonucleotide previously shown to inhibit not only cGAS but also other DNA-sensors, namely AIM2 and TLR9^21^. We observed some efficacy *in vitro* but did not see any additive or synergistic effects *in vivo* when co-loaded into NOA-pDNA-LNPs (**Supplementary Fig. 9**).

Most importantly, NOA-pDNA-LNPs completely prevented mortality in C57BL/6 mice compared to standard pDNA-LNPs at the 1 mg/kg dose (**Fig. 4l**). Notably, the mice still experience weight loss, indicating further improvements are required prior to clinical translation of NOA-pDNA-LNPs (**Supplementary Fig. 10**).

### NOA-pDNA-LNPs show prolonged transgene expression *in vivo*

We next investigated how the addition of NOA into pDNA-LNPs affects pDNA transgene expression. The duration of luciferase expression within the whole body of BALB/c mice was monitored following IV injection of either 25 μg of pDNA-LNPs or NOA-pDNA-LNPs using In Vivo Imaging System (IVIS) (**Fig. 5a**). Most importantly, pDNA-LNP and NOA-pDNA-LNP show similar levels of protein expression confirming that addition of NOA provides better safety profiles without hindering transgene expression *in vivo* (**Fig. 5b**). Notably, during this study, two mice that received 25 μg of pDNA-LNPs without NOA died within 2-days.

Furthermore, we IV injected 5 μg of mRNA-LNPs as a control and observed high luciferase protein expression that decreased back to baseline levels in ∼2 weeks (**Supplementary Fig. 11**). Note that 5 μg of mRNA approximately matches with 25 μg of pDNA in mole dose. Meanwhile, mice injected with NOA-pDNA-LNPs had significantly higher levels of luciferase protein even 32-days after LNP dose compared to PBS and mRNA-LNP controls (**Fig. 5c**).

### A small LNP formulation screen significantly boosts transgene expression of NOA-pDNA-LNPs *in vitro*

As a proof-of-concept, we investigated if we could improve protein expression from NOA-pDNA-LNPs. Typically, LNP formulation parameters - such as type and amount of ionizable lipid and helper lipid – are optimized using a screening process to improve transgene expression. Unlike mRNA delivery, there are additional challenges for pDNA delivery such as nuclear translocation which is required for pDNA transcription and pDNA degradation due to endosomal and cytosolic DNases^22,23^. As such, many studies optimizing LNP formulation for mRNA delivery may not directly translate to improving pDNA delivery.

Thus, we first performed a small Design of Experiments (DoE) screen to find optimal formulation parameters for improving the transgene expression capacity of NOA-pDNA-LNPs. Using JMP software, we designed a full factorial DoE by varying 3 parameters: amount of ALC-0315 (30-50 mole %), total lipids-to-pDNA ratio (10:1 to 40:1, w/w), and the type of helper lipid (DSPC, DOPE, DOTAP, 18:0 PG). All formulations contained NOA-to-total lipids ratio of 0.2 (mole-to-mole) and used a plasmid encoding luciferase protein. A total of 48 LNP formulations were tested in RAW264.7 cells at a dose of 500 ng/mL for 24-hours prior to measuring luciferase protein expression. DoE results show how slight changes in the LNP formulation can affect pDNA transgene expression capacity (**Fig. 6a**). LNPs formulated with DOPE or 18:0 PG as the helper lipid had lower transgene expression compared to LNPs made with DSPC or DOTAP. Importantly, optimized LNPs with DOTAP as the helper lipid led to a 50-fold increase in pDNA transgene expression (**Fig. 6b**). Optimized NOA-pDNA-LNPs were made using 40% ALC-0315, 46.4% cholesterol, 12.1% DOTAP, 1.5% ALC-0519. We speculate that DOTAP can help condense and potentially protect pDNA from cytosolic degradation leading to greater transgene expression, though further studies are required to elucidate the specific mechanisms.

To validate and generalize the optimized NOA-pDNA-LNP formulation to other cell types, we treated difficult-to-transfect cells, human induced-pluripotent-stem-cell-derived alveolar epithelial type II-like cells (iAT2s) and measured eGFP transgene expression over time. Representative images show improved transfection efficiencies of optimized NOA-pDNA-LNPs compared to the gold standard, Lipofectamine 2000 (**Fig. 6c**). Note that the tdTomato (red) signal indicates iAT2 cell positivity. 120-hours after treatment, iAT2s were collected and flow cytometry was performed which showed similar (but trending higher) % eGFP positivity in cells treated with optimized NOA-pDNA-LNPs compared to the cells treated with Lipofectamine 2000 (**Fig. 6d**). Furthermore, optimized NOA-pDNA-LNPs also showed ability to transfect precision cut lung slices (PCLS) derived from human lungs (**Supplementary Fig. 12**).

In conclusion, we demonstrate that NOA-pDNA-LNPs’ formulation can be iteratively optimized for specific applications, like improving *in vitro* transfection in difficult-to-transfect cells as shown here.

## Discussion

Despite the success of nucleoside-modified mRNA-LNP therapeutics, the relatively short half-life of mRNA expression remains one of the biggest challenges limiting its application in the treatment of chronic diseases. In contrast, pDNA delivery shows great promise, with prolonged gene expression (months) and tunable promoters providing cell-specificity and temporal control. However, here we show the acute toxicities associated with pDNA delivery via LNPs, triggering a high level of morbidity and even mortality at commonly used therapeutic doses in wild type C57BL/6 and BALB/c mice.

Through immunofluorescence staining *in vitro* and genetic knockout mice *in vivo*, we identify STING activation as a primary driver of pDNA-LNP-induced adverse events. The acute response to pDNA-LNPs is highlighted by the rapid onset of lethargy and lack of movement in mice within hours after LNP administration, culminating in 100% mortality within 2-days at 1 mg/kg dose. Furthermore, we show that this acute-but-transient inflammatory response occurs across various pDNA-LNP formulations with varying lipid and pDNA components.

To increase the safety profile of pDNA-LNPs, we introduce a novel platform technology: co-loading of endogenous anti-inflammatory lipids with STING inhibitory activity into standard pDNA-LNP formulations. We show that various AILs can be loaded into standard pDNA-LNPs without negatively affecting pDNA loading or particle stability. Specifically, we demonstrate the efficacy of nitro-oleic acid in mitigating pDNA-induced inflammation both *in vitro* and *in vivo*. By incorporating NOA into pDNA-LNPs, we achieve a significant reduction in pro-inflammatory cytokine levels and adverse events, without compromising transgene expression efficiency.

Moreover, NOA-pDNA-LNPs express transgene for at least 1 month compared to ∼2 weeks for mRNA-LNPs. We also show that NOA-pDNA-LNP formulation can be optimized for improving expression by performing a Design of Experiments (DoE) screen, revealing critical LNP formulation parameters on pDNA expression capacity. Optimized NOA-pDNA-LNPs with DOTAP as the helper lipid exhibit superior transfection efficiency compared to standard NOA-pDNA-LNPs (50-fold increase) in RAW264.7 cells and similar efficiencies to the gold-standard, Lipofectamine, even in human induced-pluripotent-stem-cell-derived type II alveolar epithelial cells (iAT2s) that are known to be difficult-to-transfect. Mechanistically, this increase in expression warrants further investigation, though we speculate it is likely due to improved protection and or condensation of pDNA.

In conclusion, our study highlights the need to address the adverse events associated with pDNA-LNP administration and presents a transformative approach to enhance the safety and efficacy of non-viral pDNA delivery. Co-loading pDNA-LNPs with bioactive molecules, such as NOA, will enable pDNA-LNPs to attain a useful position in the genetic medicine toolbox, working alongside the other clinically validated tools of mRNA-LNPs, siRNA, CRISPR, and adeno-associated viruses.

## Methods

### Materials

Ionizable lipids (D-Lin-MC3-DMA, ALC-0315, and SM-102), ALC-0159, and nitro-oleic acid (NOA) were purchased from Echelon Biosciences (Cat# N-1282, N-1102, N-1020, L-0112, respectively). Docosahexaenoic acid (DHA) was purchased from MedChemExpress (Cat# HY-B2167). Eicosapentaenoic acid (EPA) and nitro-linoleic acid (NLA) were purchased from Cayman Chemical (Cat# 90110, 30160, respectively). 18:0 PC (DSPC, 1,2-distearoyl-sn-glycero-3-phosphocholine) and DMG-PEG 2000 (1,2-dimyristoyl-rac-glycero-3-methoxypolyethylene glycol-2000) were purchased from Avanti Polar Lipids (Cat# 850365, 880151, respectively). Plasmid DNA was purchased from Aldevron (<0.1 EU/μl endotoxin level, Cat# 5078-5). 5moU nucleoside-modified firefly luciferase mRNA was purchased from TriLink BioTechnologies (Cat# L-7202).

### Animals

All animal experiments strictly adhered to the guidelines established in the Guide for the Care and Use of Laboratory Animals (National Institutes of Health, Bethesda, MD). Approval for all animal procedures was obtained from the University of Pennsylvania Institutional Animal Care and Use Committee. naïve C57BL/6, naïve BALB/c, and STING-knockout BALB/c mice, aged 6–8 weeks and weighing 23-25 g, were procured from The Jackson Laboratory, Bar Harbor, ME, for the study. The mice were housed in a controlled environment maintained at temperatures between 22–26 °C, with a 12/12-hour light/dark cycle, and provided with access to food and water.

For survival curve studies, mice were monitored and weighed daily. Any mice with visual cues of extreme distress or weight loss >20% were euthanized and removed from the study. All intravenous injections were done retro-orbitally by injecting into the retro-bulbar sinus.

### LNP Formulation

LNPs were formulated using microfluidics (NanoAssemblr Ignite, Precision Nanosystems). Lipids were dissolved in ethanol and mixed with aqueous buffer (50 mM citrate buffer, pH 4) containing either 5moU nucleoside-modified mRNA or pDNA, at a total flow rate of 6 mL/min, a flow rate ratio of 1-to-3 (ethanol-to-aqueous), and total lipid-to-nucleic acid ratio of 40-to-1 (w/w). LNPs were dialyzed against 1x PBS in a 10 kDa molecular weight cut-off cassette for 2 h, stored at 4 °C, and used within 2-days. Anti-inflammatory lipids (AILs) were added as a 5^th^ component at a drug-to-total lipid ratio of 0.2 (mole-to-mole) for all studies using AIL loaded pDNA-LNPs unless otherwise indicated.

All LNPs made used FDA-approved formulations with the following molar ratios (D-Lin-MC3-DMA LNPs: 50% D-Lin-MC3-DMA, 38.5% cholesterol, 10% DSPC, 1.5% DMG-PEG 2000; SM-102 LNPs: 50% SM-102, 38.5% cholesterol, 10% DSPC, 1.5% DMG-PEG 2000; ALC-0315 LNPs: 46.3% ALC-0315, 42.7% cholesterol, 9.4% DSPC, 1.6% ALC-0159). All studies used ALC-0315 LNP formulation unless otherwise indicated.

### LNP Characterization

Measurements of hydrodynamic nanoparticle size and polydispersity index was conducted through dynamic light scattering (DLS) using a Zetasizer Pro ZS (Malvern Panalytical). The encapsulation efficiencies and concentrations of LNP mRNA or pDNA were determined using a Quant-iT RiboGreen RNA assay or Quant-iT PicoGreen dsDNA assay, respectively (Invitrogen). Anti-inflammatory lipid (AIL) drug loading in LNP was determined using ultra-performance liquid chromatography (UPLC, UV/Vis) after purifying out unloaded AIL using size exclusion Zeba Spin Desalting Columns (Thermo).

### Cell Culture

RAW264.7 mouse macrophages were purchased from ATCC and cultured in Dulbecco’s modified Eagle’s medium (DMEM) with 10% heat-inactivated fetal bovine serum (FBS) and 1% penicillin/streptomycin (PS). All cells were incubated with 5% CO2 at 37°C.

For cell viability and luciferase assays, cells were seeded at a density of 1E5 cells/well in a 96 (clear bottom for cell viability assay and white bottom for luciferase assay) well plate using 100 uL 24h prior to LNP treatment. cck8 assay was performed according to manufacturer’s instructions to measure cell viability % (Abcam). “Luciferase Assay Systems” protocol was used to measure expression capacity of either mRNA or pDNA LNPs *in vitro* (Promega).

For cytokine studies, cells were seeded at a density of 3E5 cells/well in a 24 clear bottom well plate using 300 uL 24h prior to LNP treatment. 4-hours after LNP treatment, supernatant was collected, spun down at 10,000 g for 10 min to remove any debris and stored at -80°C.

For immunofluorescence imaging studies, cells were seeded at a density of 1.2E5 cells/well in 8-well μ-Slide chambers.

### p-STING and p-TBK1 imaging and quantification

Cells were treated with LNPs for 4-hours. Then, they were fixed in 4% fresh paraformaldehyde for 15 min. Permeability was then performed with 0.05% saponin buffer (J63209.AK, Invitrogen) for 10 min. Then, samples were incubated with 10% normal goat serum (50062Z, Life Technologies Corp.) for 1 h. After PBS washing, cells were treated with 1:150 primary antibodies (anti p-STING, 62912S, Cell Signaling Technology Inc.; anti p-TBK1, 5483S, Cell Signaling Technology Inc.) at 4°Covernight. Cells were washed with PBS to remove unbound antibodies. Next, cells were incubated with 1:750 secondary antibodies (Alexa Fluor 488-conjugated goat anti-rabbit antibody, A11008, Invitrogen) at 37 ^°^C for 1.5 h. Cell nuclei were labeled using DAPI. Images were acquired by LSM980 microscopy (Zeiss) and mean fluorescent intensity (MFI) was measured by Image J.

### iPSC-derived Alveolar type 2 epithelial cell transfection

Alveolar type 2 epithelial cells derived from human induced pluripotent stem cells (iAT2s) were maintained in 3D growth factor reduced (GFR)-Matrigel culture as previously described^24,25^. Plating iAT2s on 2D transwell inserts (6.5mm; Falcon) was performed as previously described^26^. In brief, transwell inserts were coated with diluted hESC-Qualified Matrigel (Corning) as instructed by the manufacturer. A single-cell suspension of iAT2s from 3D culture was obtained by dissociating Matrigel droplets for 30 minutes with 2 mg/mL Dispase (Gibco) followed by 15 minutes of 0.05% trypsin-EDTA (Gibco) at 37 °C. iAT2s were plated on pre-coated transwell inserts at a density of 500,000 live cells/cm^2^ in 500 μL of CK+DCI+Y (3 μM CHIR99021, 10 ng/mL KGF, 50 nM dexamethasone, 0.1 mM cAMP, 0.1 mM IBMX, 10 μM Rho-associated kinase inhibitor (Y) [MilliporeSigma, Y-27632]), with 500 μL of CK+DCI+Y added to the basolateral compartment. 48-hours after plating, iAT2s were refreshed with 500uL CK+DCI in both the apical and basolateral compartments prior to LNP administration and transfection.

Various conditions of LNPs containing eGFP pDNA were administered dropwise onto 2D cultures of iAT2s. Lipofectamine 2000 (Thermo) transfection of eGFP pDNA was performed according to the manufacturer’s protocol. iAT2s were imaged for tdTomato retention and eGFP expression using an Eclipse Ti2 Series inverted microscope (Nikon) and 24-, 48-, and 120-hours after treatment. After 120 hours, iAT2s were isolated from transwell inserts with Accutase (StemCell) and washed in FACS Buffer (0.1% BSA, 2mM EDTA, in PBS pH 7.4) for flow cytometry analysis. iAT2 tdTomato retention and eGFP expression was assessed by flow cytometry using a CytoFlex SRT (Beckman) and analyzed using FlowJo v10.10 software.

### Transfection of precision cut lung slices (PCLS) from human lung tissues

Human PCLS of 300 μm were cultured in DMEM with 10% FBS and 1% Primocin overnight at 37°C. Transfection mixture of plasmid eGFP (1000 ng/mL) was mixed with transfection reagent Lipofectamine 2000 (Cat# 11668) in Opti-MEM before adding to PCLS. Optimized NOA-pDNA-LNPs (1000 ng/mL) were prepared in DMEM with 10% FBS and 1% Primocin. Mixtures were added in PCLS cultured in DMEM with 10% FBS and 1% Primocin and incubated overnight at 37°C. 24-hours after transfection, eGFP tissue imaging was performed. Images were taken using Leica confocal microscope. The aspect ratio of these images was reduced 3:5 followed by 4:5, 2 times to highlight area of interest.

### In Vivo Plasma Collection

Mice treated with LNPs were sacrificed by terminal blood collection via inferior vena cava. Opening of the major body cavity and subsequent thoracotomy was performed as a secondary measure of sacrifice. Blood was centrifuged at 1000 g for 10 mins (room temperature), then plasma supernatant was collected and stored at -80°C.

### *In Vitro* and *In Vivo* Cytokine Measurements

Cytokine measurements were carried out on plasma (2x diluted) or cell culture supernatant (undiluted) with a LegendPlex 13-plex Mouse Inflammation Panel (Biolegend) according to the manufacturer’s instructions.

### In Vivo Imaging System (IVIS)

At the time of imaging, mice were intraperitoneally injected with 100 uL of 30 mg/mL D-luciferin sodium salt (Regis Technologies Inc, 103404-75-7) under 3% isoflurane-induced anesthesia, then placed in an IVIS Spectrum machine (PerkinElmer) belly up and imaged for whole body chemiluminescence every 0.2 minutes with automatically determined exposure time for 10-12 images, until the signal reached the peak intensity.

### Statistics

All results are expressed as mean ± SEM unless specified otherwise. Statistical analyses were performed using GraphPad Prism 8 (GraphPad Software) * denotes p<0.05, ** denotes p<0.01, *** denotes p<0.001, **** denotes p<0.0001.

## Acknowledgments

We thank the laboratories of Drs. Darrell Kotton and Konstantinos-Dionysios Alysandroatos for gifting Alveolar type 2 epithelial cells derived from human induced pluripotent stem cells (iAT2s).

Research reported in this publication was supported by the American Heart Association under Grant 24PRE1195406 (to M.N.P), Grants R01-HL-153510, R01-HL160694, R01-HL164594 (to J.S.B), and Grant R01AI153064 (to N.P).

