## Supplementary Figures for "Enabling non-viral DNA delivery using lipid nanoparticles co-loaded with endogenous anti-inflammatory lipids"

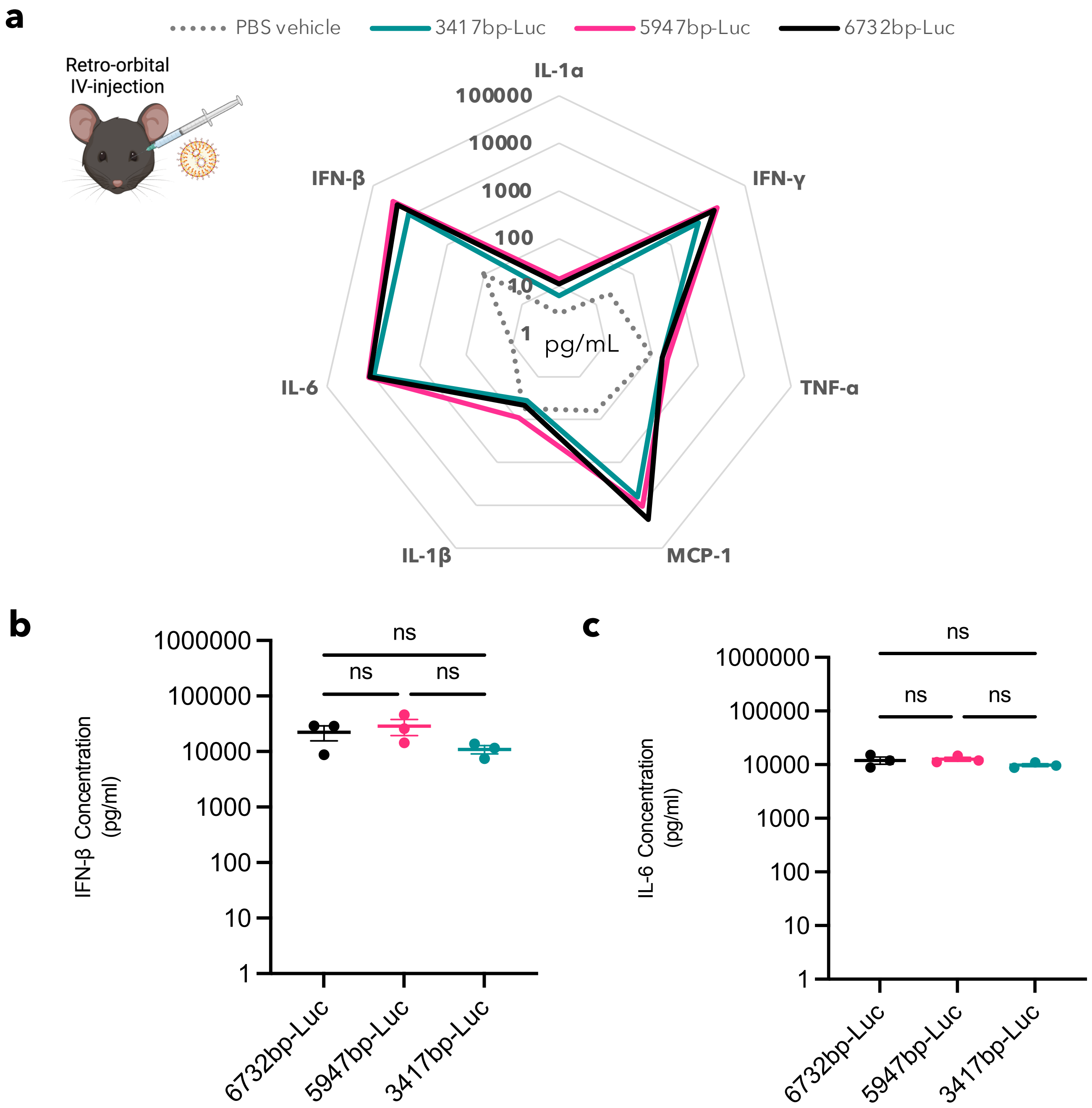


**Supplementary Figure 1: pDNA-LNPs induce acute inflammation regardless of the plasmid backbone used.**

**a, b, c,** Multiplex analysis of pro-inflammatory plasma cytokines 4-hours post 5 µg IV dose of pDNA-LNPs indicates acute systemic inflammation regardless of plasmid size with no significant change in IFN-β (**b**) and IL-6 **(c**) levels.

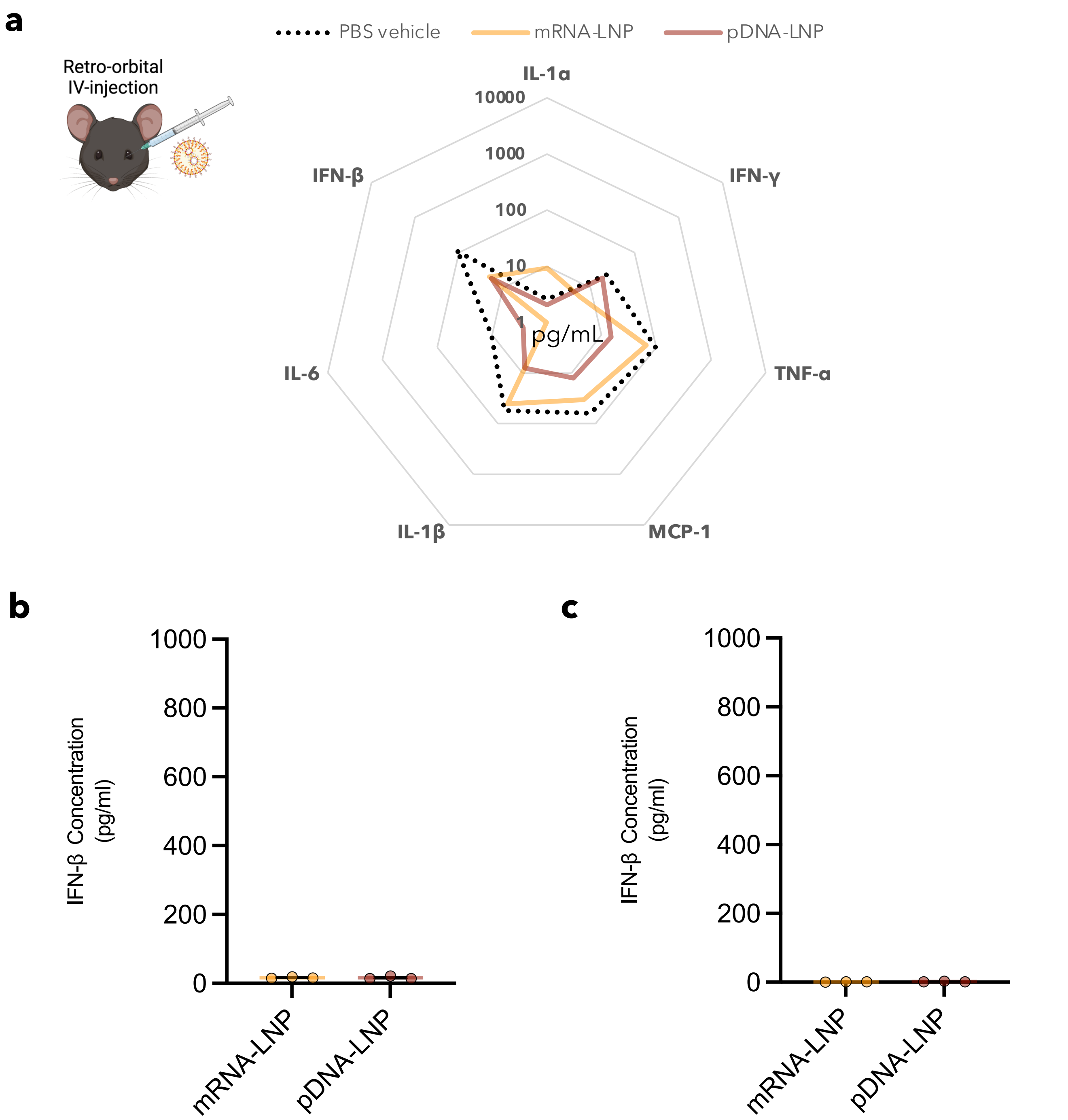


**Supplementary Figure 2: pDNA-LNP-induced inflammation does not reappear at later timepoint**

**a, b, c,** Multiplex analysis of plasma cytokines 5-days after 5 μg IV injection of mRNA- or pDNA-LNPs shows all cytokines levels are back to baseline levels, specifically IFN-β (**b**) and IL-6 (**c**).

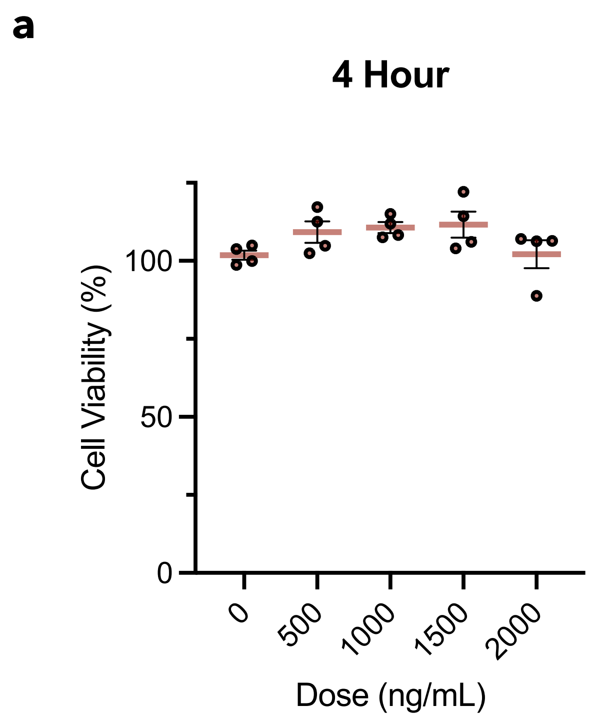


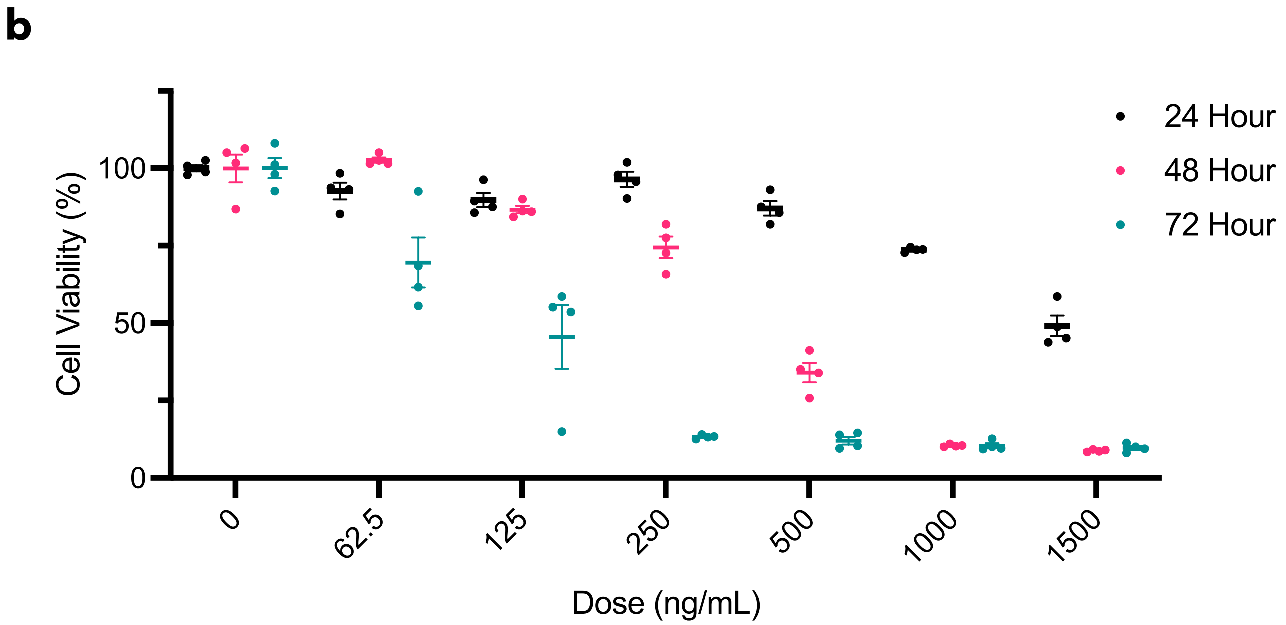


**Supplementary Figure 3: Cell viability after pDNA-LNP treatment in RAW264.7 cells.** **a,** cell viability measured 4-hours after various doses of pDNA-LNPs shows good tolerability. Note that all studies measuring cell supernatant cytokines were done 4-hours after 1000 ng/mL dose where cell viability is ~100%. **b**, Cell viability as a function of dose- and time-response of pDNA-LNPs.

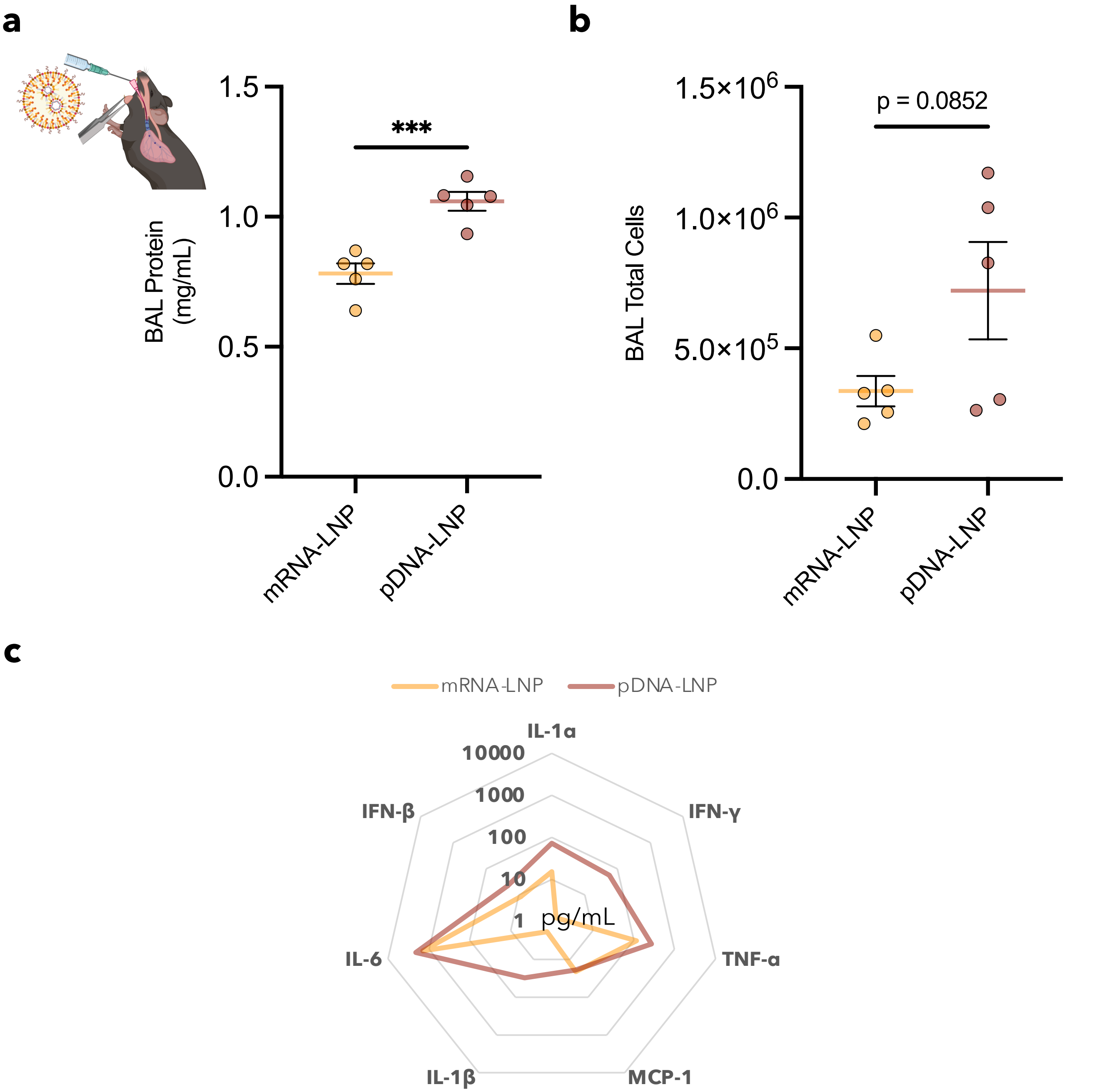


**Supplementary Figure 4: pDNA-LNPs also induce acute inflammation when administered intratracheally in naïve C57BL/6 mice.**

**a, b, c,** After intratracheally administering 5 μg mRNA- or pDNA-LNPs, we examined inflammation specific to the lungs by examining protein and leukocyte levels in the bronchoalveolar lavage (BAL) fluid, which indicates capillary leakage and leukocyte penetration into the alveoli (air sacs). Protein (**a**), total cells (**b**), and pro-inflammatory cytokines (**c**) were higher in the BAL fluid from mice that were administered pDNA-LNPs compared to the ones given mRNA-LNPs.

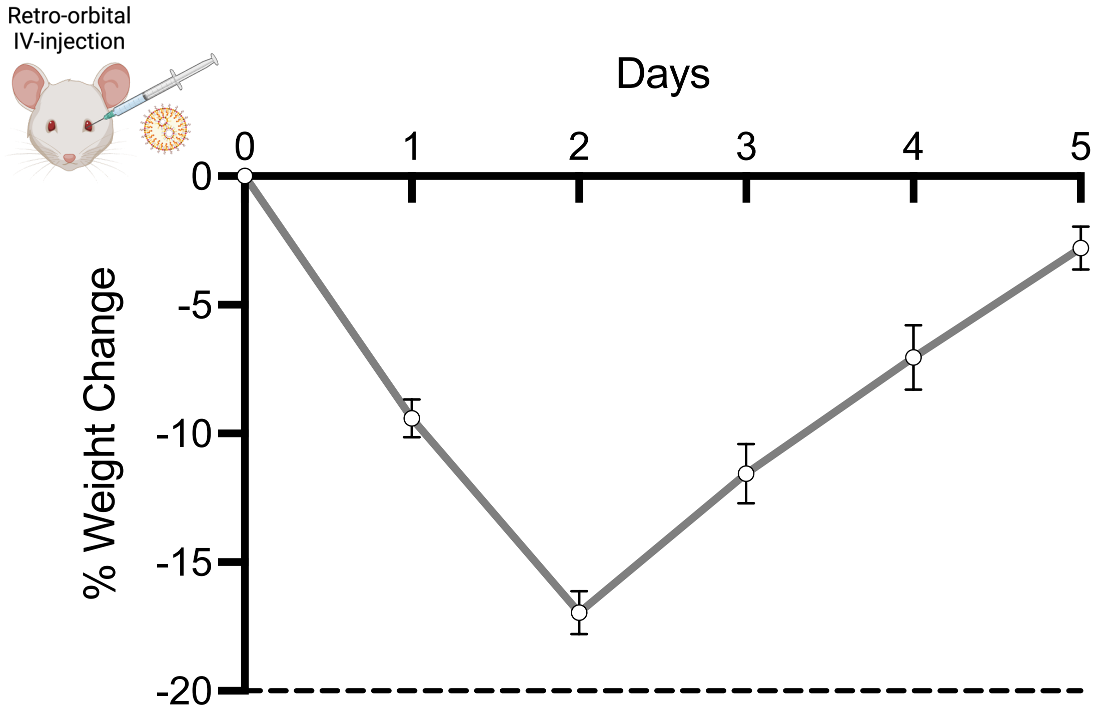


**Supplementary Figure 5: pDNA-LNPs cause extreme weight loss in naïve BALB/c mice.**

Weight was monitored over time of naive BALB/c mice that survived after IV-injection of 1 mg/kg (~25 µg) of pDNA-LNPs.

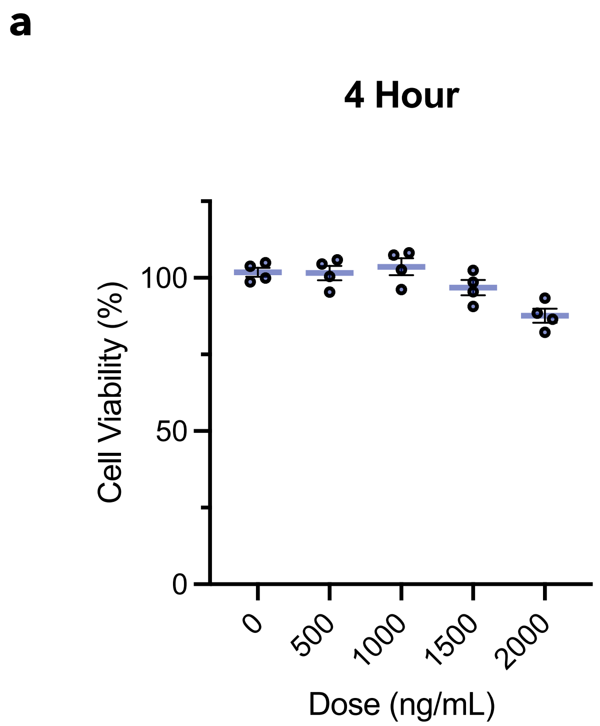


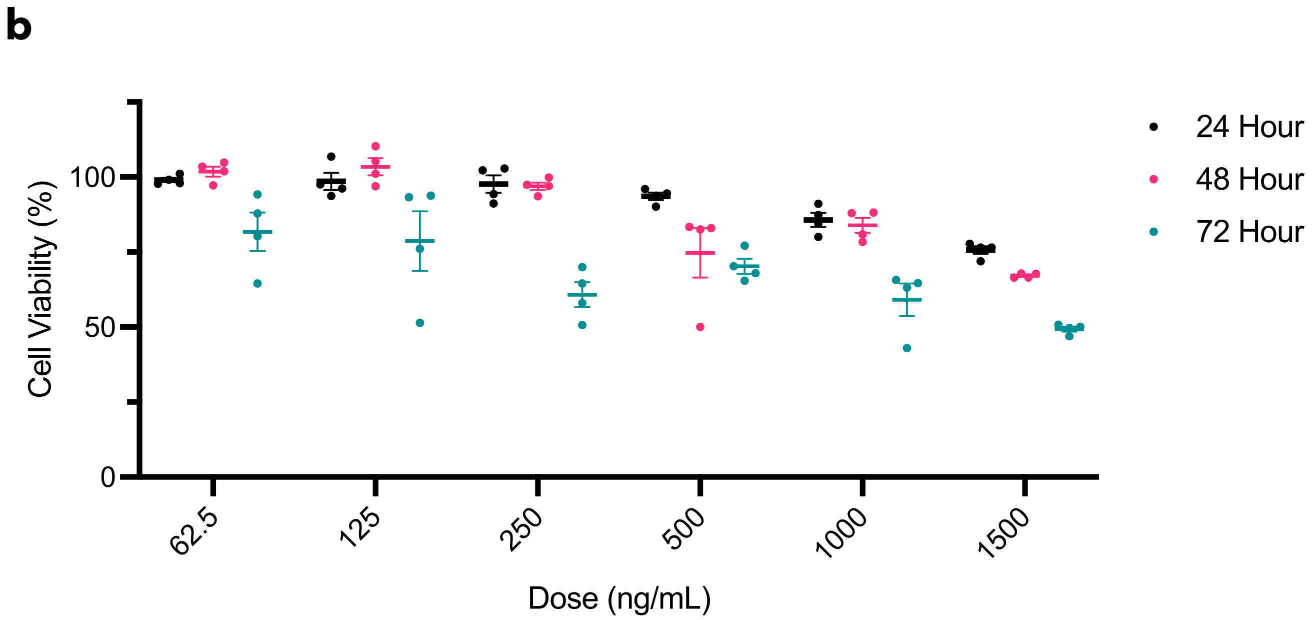


**Supplementary Figure 6: Cell viability after NOA-pDNA-LNP treatment in RAW264.7 cells.**
**a,** cell viability measured 4-hours after various doses of NOA-pDNA-LNPs shows good tolerability. Note that all studies measuring cell supernatant cytokines were done 4-hours after 1000 ng/mL dose where cell viability is ~100%. **b**, Cell viability as a function of dose- and time-response of NOA-pDNA-LNPs.

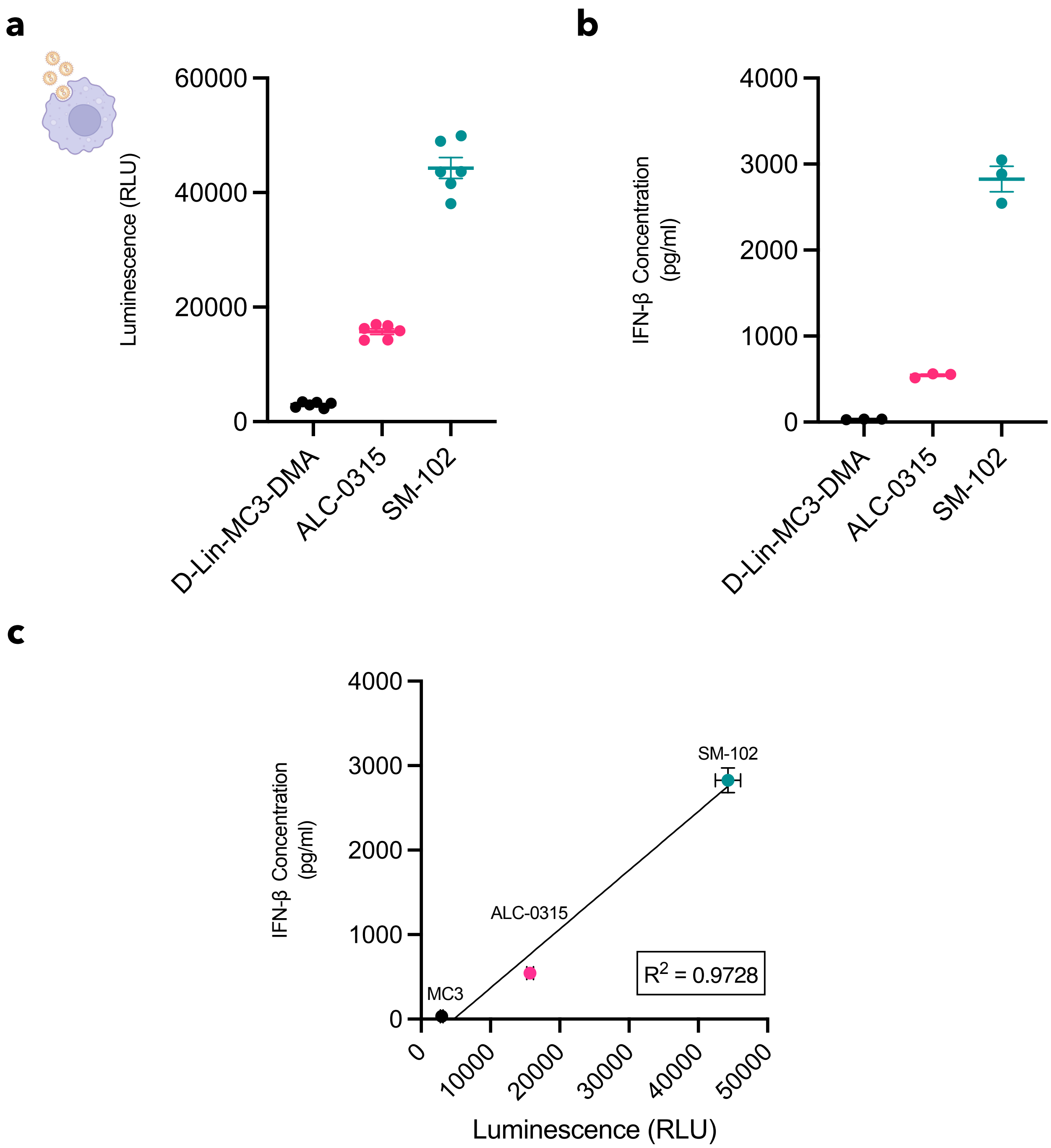


**Supplementary Figure 7: Transfection efficiencies of pDNA-LNPs correlate with the levels of IFN-b secretion in RAW264.7 cells.**

**a, b, c,** RAW264.7 cells were incubated with 1000 ng/mL of pDNA-LNPs made from 3 FDA-approved formulations (Patisiran [D-Lin-MC3-DMA], mRNA-1273 [SM-102], and BNT162b2 [ALC-0315]). pDNA-LNPs that leads to higher transgene expression (**a**) leads to greater levels of IFN-β (**b**) showing a linear correlation for the 3 FDA formulations tested (**c**).

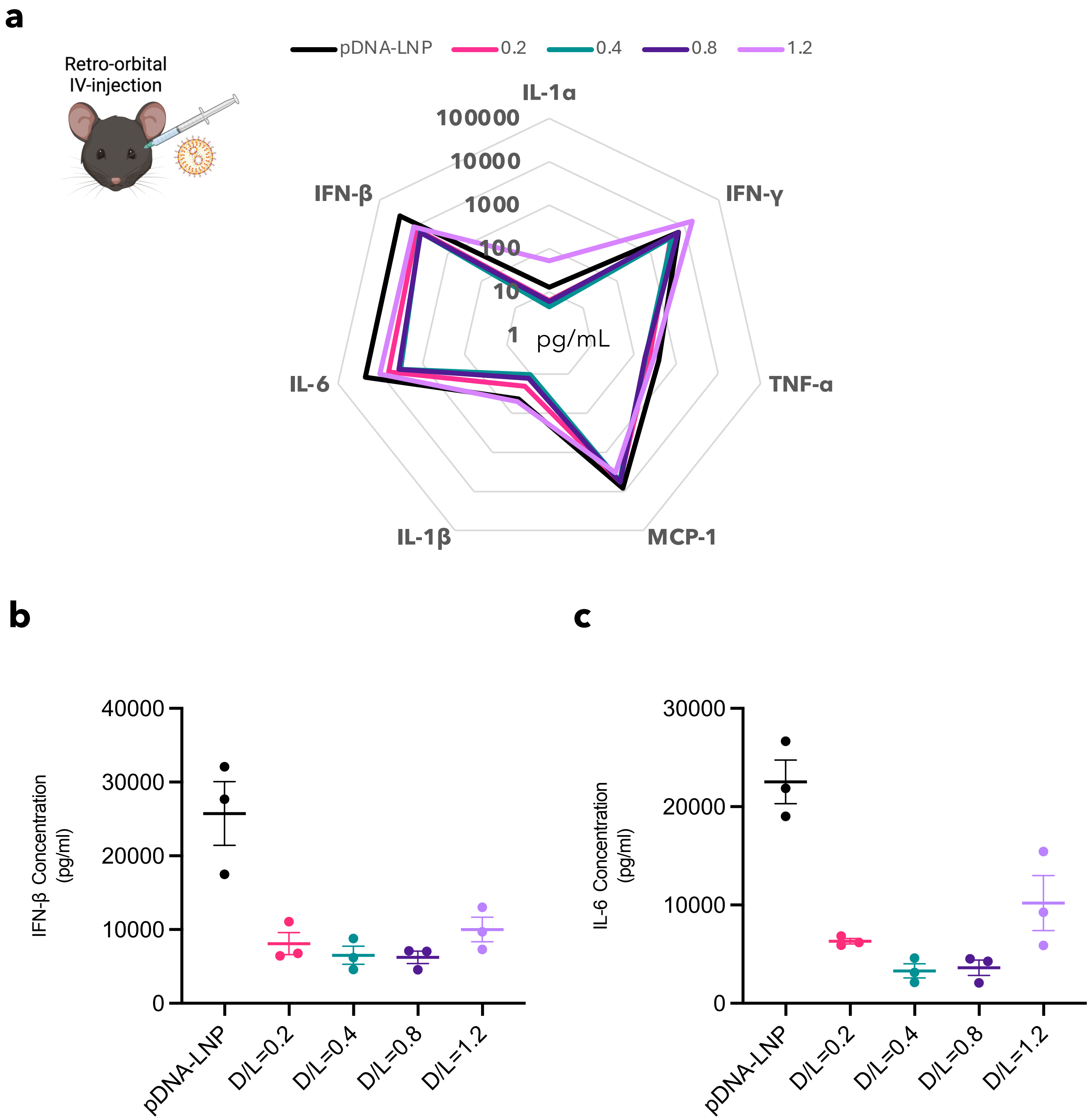


**Supplementary Figure 8: Dose-response of NOA in NOA-pDNA-LNPs**
**a, b, c,** 5 µg of pDNA-LNPs were formulated with 0.2-1.2 NOA-to-total lipid ratio (mole-to-mole) were IV injected into naïve mice. Multiplex analysis of pro-inflammatory plasma cytokines 4-hours after 5 µg IV-injection indicates 0.2-0.8 NOA-to-total lipid ratio as the optimal ratio for the best reduction of most pro-inflammatory cytokines (**a**), specifically IFN-β (**b**) and IL-6 (**c**).

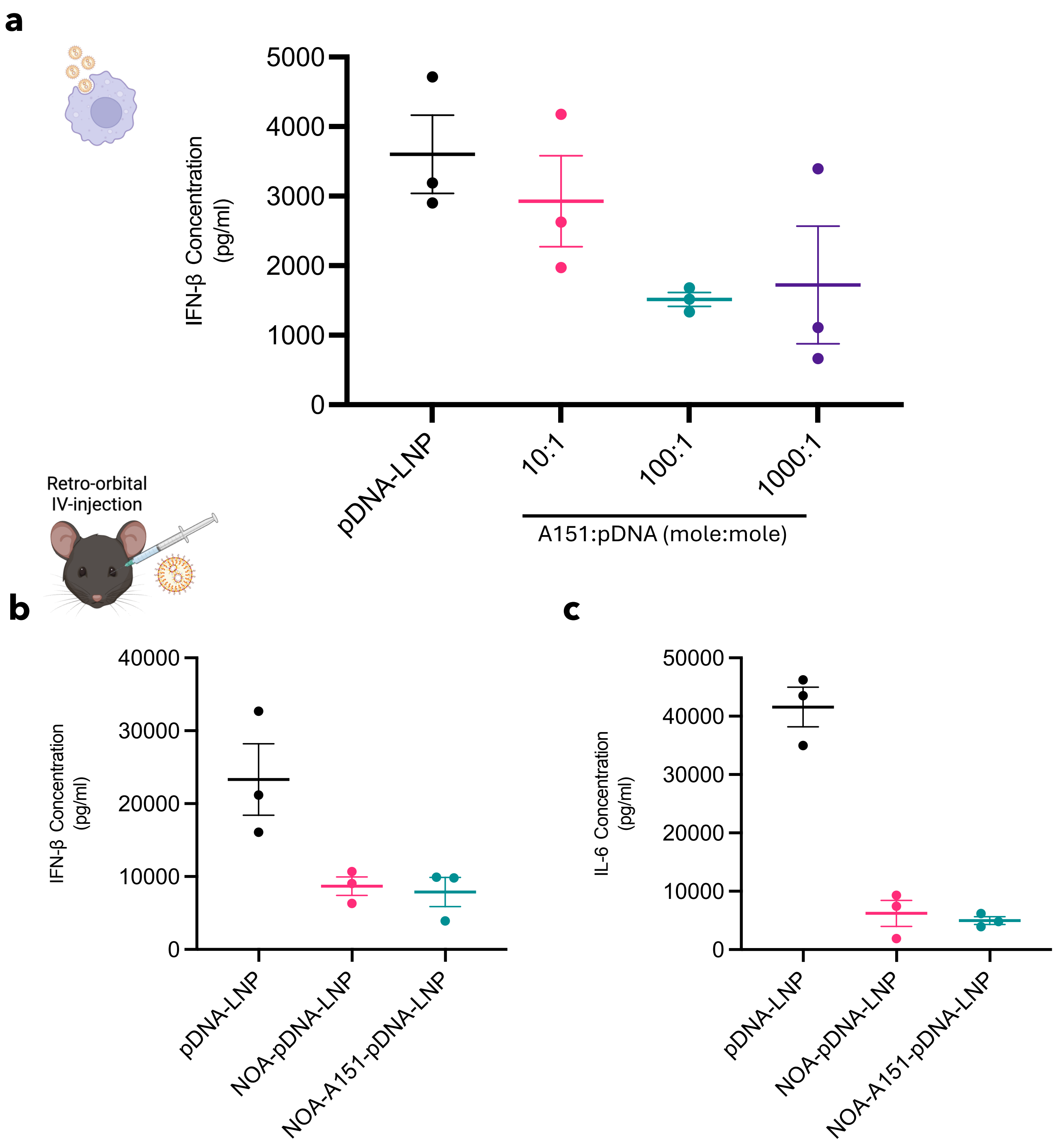


**Supplementary Figure 9: A151, oligonucleotide inhibitor of not only cGAS but also other DNA sensors (namely AIM2 and TLR9), reduces pDNA-LNP inflammation only *in vitro***

**a,** Dose-response of A151-loaded pDNA-LNPs show reduction of IFN-β release in RAW264.7 cells. **b, c**, Co-loaded A151 with NOA-pDNA-LNPs does not have an additive or synergistic effect of reducing pDNA-induced inflammation in naïve mice.

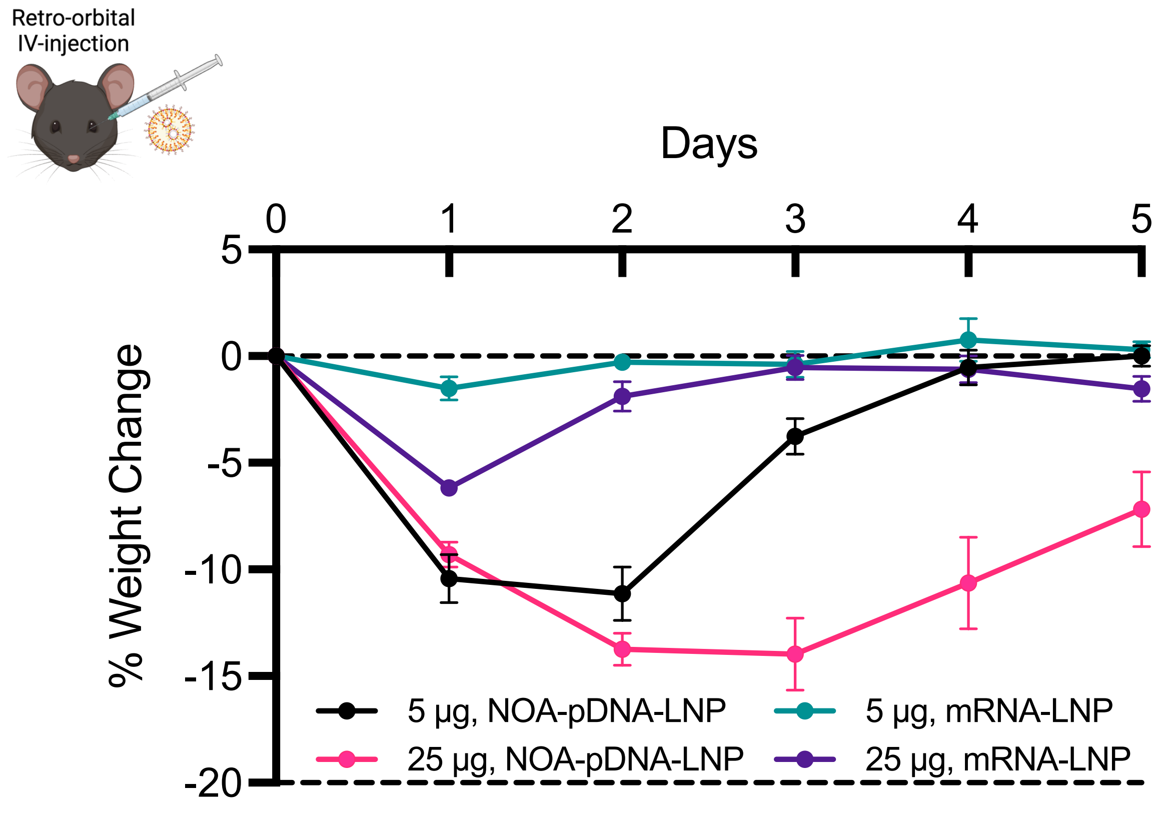


**Supplementary Figure 10: Weight loss of C57BL/6 mice treated with mRNA-LNP and NOA-pDNA-LNPs**

Weight change over time for Naïve C57BL/6 mice that were injected with either 5 µg or 25 µg of mRNA-LNPs or NOA-pDNA-LNPs.

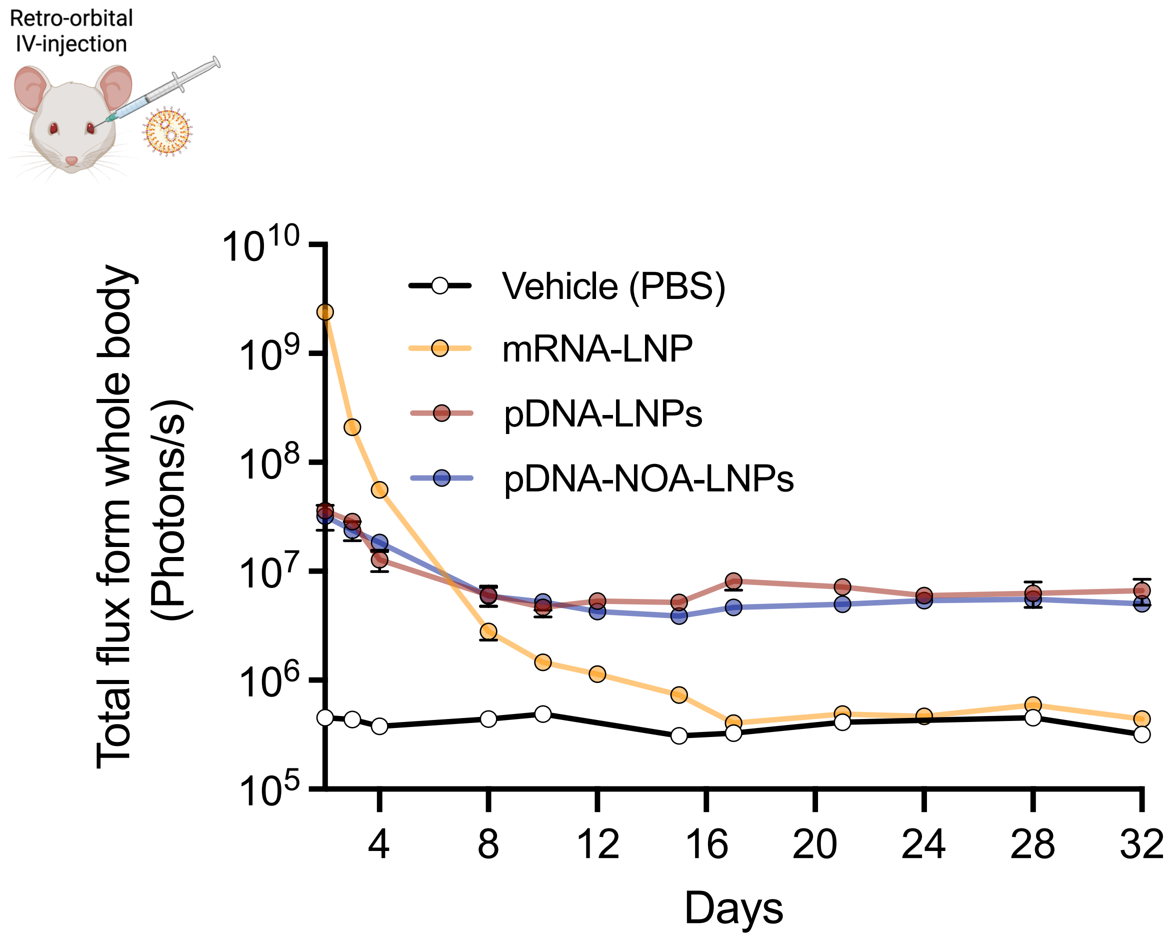


**Supplementary Figure 11: mRNA-LNP control for IVIS imaging**
Quantified total flux (photons/s) from IVIS images from Naïve BALB/c mice IV injected with 5 µg of 5moU nucleoside modified mRNA-LNPs, 25 µg of pDNA-LNPs, or 25 µg of NOA-pDNA-LNPs. Note that 5 µg of mRNA approximately matches 25 µg of pDNA in mole amount.

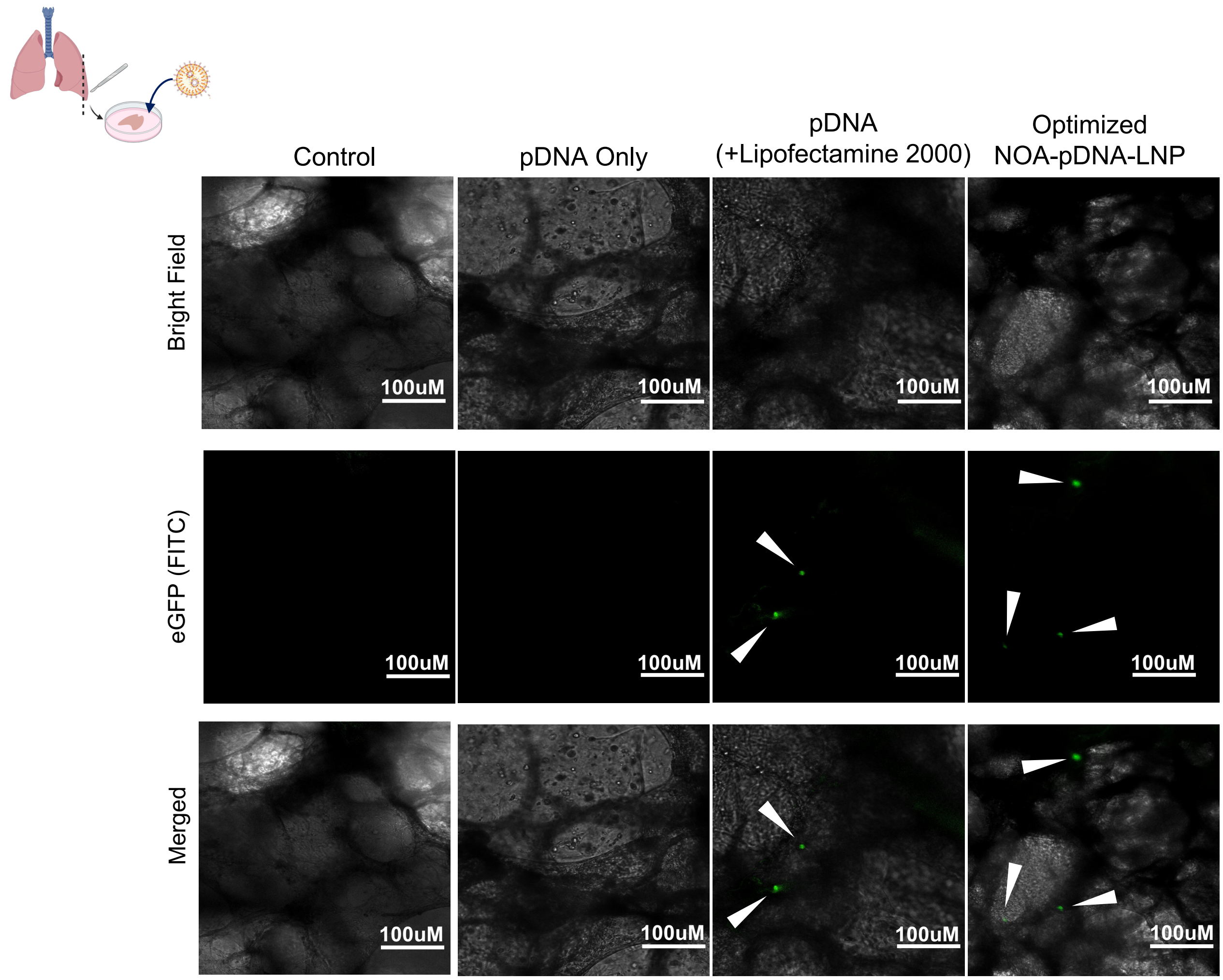


**Supplementary Figure 12: pDNA transfection in precision cut lung slices (PCLS) from human lung tissue**

Precision cut lung slices from human lung tissue were plated and cultured. eGFP pDNA was transfected with either Lipofectamine 2000 or optimized NOA-pDNA-LNPs at a dose of 1000 ng/mL. 24-hours later, eGFP images were captured showing similar levels of transfection for PCLS treated with Lipofectamine 2000 and optimized NOA-pDNA-LNPs.
